# A shift from competition to facilitation amplifies the temperature-dependence of microbial community respiration

**DOI:** 10.1101/2021.04.15.439947

**Authors:** Francisca C. García, Tom Clegg, Daniel Barrios O’Neill, Ruth Warfield, Samraat Pawar, Gabriel Yvon-Durocher

## Abstract

The respiratory release of CO_2_ by microbes is a dominant component of the global carbon cycle. However, large uncertainties exist about the effects of climatic warming on the respiration of microbial communities due to lack of mechanistic, empirically-tested theory that accounts for dynamic species interactions. We developed a general mathematical model which predicts that thermal sensitivity of microbial community respiration increases as species interactions become more positive, i.e., change from competition to facilitation. This is because facilitation disproportionately increases positive feedbacks between the thermal sensitivities of species-level metabolic and biomass accumulation rates at warmer temperatures. We experimentally validated this prediction in bacterial communities of 8 taxa, finding that a shift from competition to facilitation after a month of co-adaptation caused a 60% increase in the thermal sensitivity of their respiration relative to *de novo* communities that had not co-adapted. Thus, rapid changes in species interactions can profoundly change the temperature-dependence of microbial community respiration and should be considered in climate change models.

## MAIN

Empirical data show that ecosystem-level respiration generally follows an exponential-like relationship with temperature^1^ These findings have led to concerns that climatic warming will increase carbon emissions from the biosphere, increasing positive feedbacks in the carbon cycle, ultimately accelerating the rate of planetary warming^2–5^. Microbes, and in particular bacteria, by conservative estimates make up ~20% of earth’s total biomass^6^, and by decomposing organic matter, account for a major fraction of the thermal response of ecosystem-level respiration^2,7^. For example, bacterial contribution to ecosystem respiration is estimated to be >50% in some ocean biomes^7,8^. Consequently, even small changes in the thermal sensitivity of microbial community respiration will likely have significant impacts on future global warming projections^8,9^. However, the response of microbial community respiration to temperature changes remains a key uncertainty in climate-carbon cycle projections for the coming century, and is also an unresolved question in microbial ecology^2,10,11^.

Published models of temperature responses of complex ecosystems typically assume that thermal responses can be scaled up from individual- to the ecosystem-level by a simple, weighted sum of the temperature responses of component species’ populations^9,12–15^. These models only focus on the direct effect of temperature on individuals and species’ metabolism, ignoring the effects of interactions among species. However, species interactions such as predation, competition and facilitation drive population dynamics in all ecosystems, determining the amount and distribution of biomass across species’ populations, and ultimately total ecosystem respiration and its response to changes in temperature^16,17–21^. Thus, by failing to account for the effects of species interactions, current models may not be able to predict the response of ecosystem respiration to changing temperatures.

In microbial communities in particular, demographic processes and population turnover occur over relatively short timescales, and the temperature-dependence of community respiration most likely reflects the direct effect of temperature on individual metabolism as well as its indirect effects through species interaction-driven biomass dynamics. Microbial taxa interact in numerous ways, ranging from competition for limiting abiotic resources, to facilitation through cross-feeding on metabolic by-products^18–20^. Microbial metabolic traits are temperature-sensitive, so when temperatures change this alters interaction-driven biomass dynamics and thus community-level respiration^21^. For example, widespread facilitation might amplify the effects of temperature by creating a positive feedback loop due to enhanced metabolic and growth rates in warmer conditions. Conversely, if weak or neutral interactions occur such as when species partition resources, their populations might become relatively decoupled, resulting in a response of community respiration to temperature that is a simple sum of the thermal responses of individual taxa weighted by their respective population biomasses. In general, if species interactions are strong, so will be the feedbacks between populations, amplifying the (positive or negative) effects of temperature across the whole system. Here, we investigated the role of biotic interactions in the temperature dependence of community metabolism by combining theory with laboratory experiments.

## RESULTS

### Modelling the temperature dependence of microbial community respiration

Our mathematical model links the effects of species-level metabolism and inter-species interactions to the thermal response of community-level (henceforth, synonymous with “ecosystem-level”) respiration (Fig 1). A fundamental premise of our model is that species interactions act primarily to affect species’ biomasses and have a negligible effect on individual-level respiration rate. Respiration rate is constrained by cellular enzyme kinetics and is therefore driven primarily by environmental temperature. We also focus on the stages of community assembly and dynamics before populations reach carrying capacity for two key reasons. First, the bulk of community respiratory flux occurs when resource availability is high e.g., spring blooms in seasonal aquatic systems^22^ and litter fall in soils^23^, during which populations are in near exponential growth. Second, environmental perturbations and immigration events in natural microbial communities mean that these communities are constantly perturbed from equilibrium over time^24,25^.

**Fig 1.**
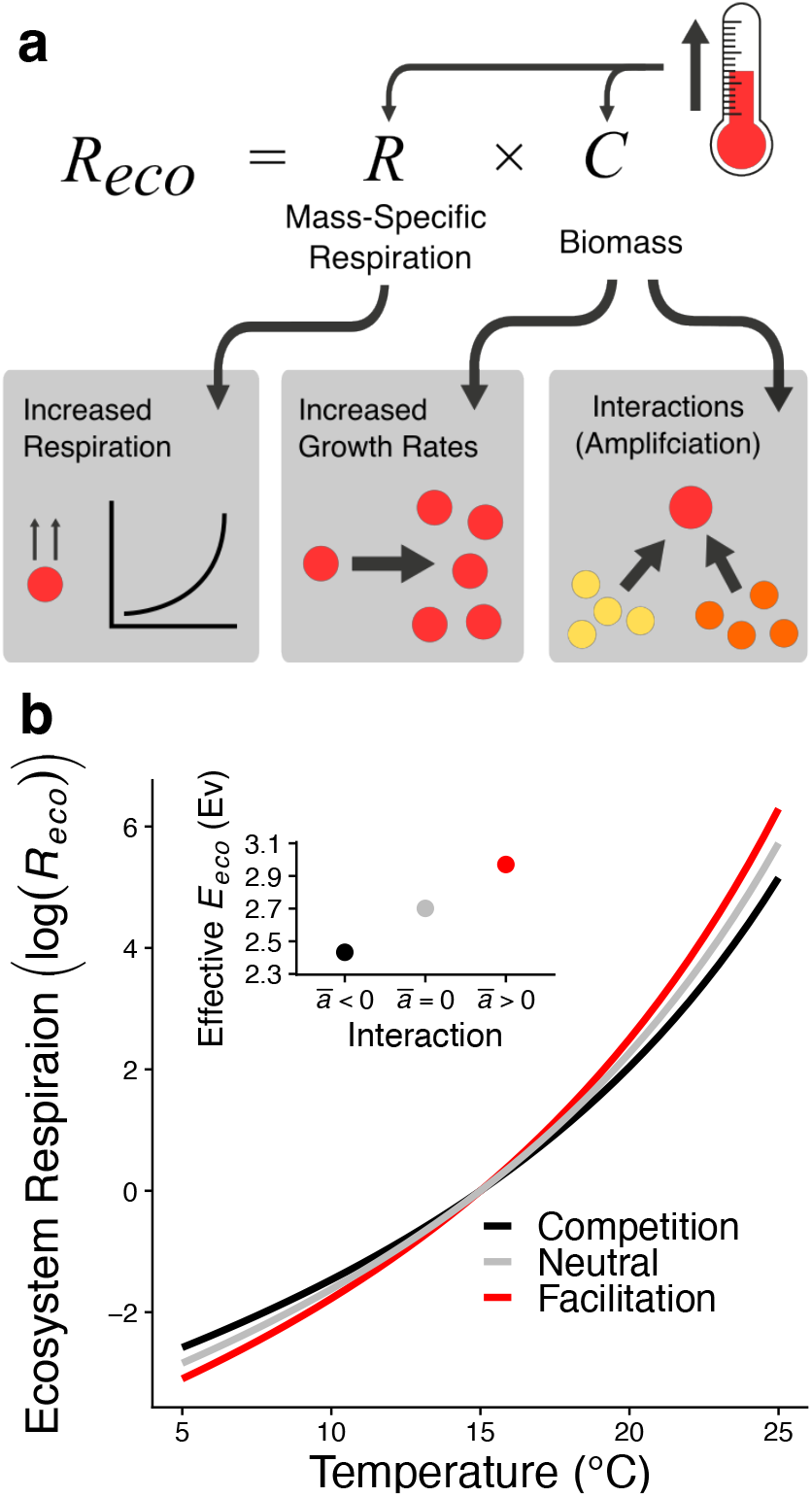
Species interactions affect the temperature sensitivity of microbial community respiration. a) Temperature can act on community-level respiration either by affecting individual metabolism directly (increasing respiration), or the amount of biomass, which is determined by the effects of temperature on growth rates and interactions between species. b) Prediction of the relationship between temperature and community respiration under different interaction structures. Respiration becomes more sensitive to temperature change as interactions become more positive. Main plot shows community respiration, log (*R_eco_*) normalised to a common *T_ref_* (15°C) at three levels of interaction strength across the community. Note that the curves are nonlinear in log scale because the thermal sensitivity of community respiration (the slope of log (*R_eco_*) versus temperature) is itself temperature-dependent (Eq 2) due to the nonlinear change in biomass dynamics with temperature as explained in the main text. Inset plot shows the resultant effective thermal sensitivity *E_eco_* measured at *T_ref_* See Methods for parameter values used to generate these specific theoretical predictions.

Consider a community comprising of *N* interacting species. This community’s total temperature (*T*)-dependent respiration rate (*R_eco_*(*T*)) depends on the sum of contributions of each population’s total respiration, which in turn can be expressed as the product of mass-specific respiration (*R_i_*(*T*)) and biomass (*C_i_*(*T*)) of each population:

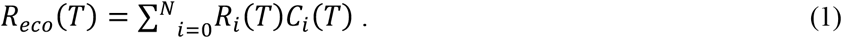

This equation implies that temperature affects community respiration by changing mass-specific respiration of individual populations, by changing their biomasses, or both (Fig 1a). Next, we derive the thermal sensitivity of *R_eco_*(*T*) (the magnitude of change of community respiration to a unit change in temperature in log-scale), which we denote by an apparent activation energy, *E_eco_* (see Methods):

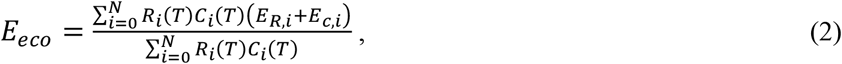

where *E_R,i_* and *E_C,i_* are the thermal sensitivities (apparent activation energies) of mass-specific respiration and biomass dynamics of the *i*^th^ species’ population, respectively. Eq 2 shows that *E_eco_* is given by the average thermal sensitivities of biomass dynamics and respiration across all species (strains) in the system, weighted by each species’ total respiratory output, *R_i_*(*T*)*C_i_*(*T*). Note that Eq 2 also contains temperature-dependent terms reflecting the effects of biomass dynamics, which results in a non-exponential thermal response of total ecosystem respiration (Fig. 1b). Next, we consider how *E_eco_* (Eq 2) is affected by pairwise interspecific interactions. Below, we will consider the potential effects of indirect and higher order interactions (see Supplementary Materials). Assuming interactions do not affect species’ mass-specific respiration rate *R_i_* (i.e., a single strain cell will have roughly the same respiration rate in the presence or absence of interactions), we focus on how they affect the biomass terms (*C_i_*’s and *E_C,i_*’s) in Eq 2. We show that the thermal sensitivity of community-level respiration can be partitioned as (see Methods)

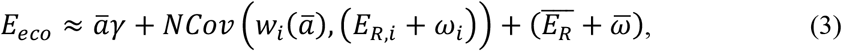

where *<u>a</u>* is the average of the interaction coefficients between all species pairs 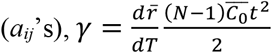 is the average thermal sensitivity of time (*t*)-dependent biomass across all populations (driven by temperature-dependent changes in average population growth rate 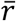), 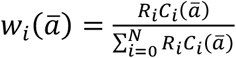 is the weighting of the *i*^th^ species’ contribution (its normalised respiratory output), and 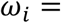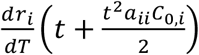 is the thermal sensitivity of its biomass at time *t*. The dependence of these weights on *ā* arises through the effects of species interactions on population biomasses (*C_i_*(*ā*)).

Equation 3 shows that the thermal sensitivity of community respiration is determined by three components: (i) the effect of averaged interspecies interactions (*<u>a</u>γ*), (ii) the covariation between species’ responses to interactions (via the weighting terms) and their thermal sensitivities (*NCov*(*w_i_*(*ā*), *E_R,i_* + *ω_i_*)) and (iii) the average thermal sensitivities of respiration and biomass growth (not accounting for interactions) across species in the community 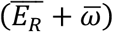. When interactions are on average neutral (*ā* = 0) the first of these terms is zero and community sensitivity is dependent only on internal factors affecting species’ population growth (growth rates and intra-specific interactions in the *ω* and covariance term). When interactions are non-neutral (*ā* ≠ 0) they will either amplify or dampen the sensitivity of community respiration relative to this neutral case (Fig 1b). More facilitatory (positive) interactions will increase (amplify) thermal sensitivity, while competitive interactions will result in a reduction (dampening) in sensitivity (including intransitive competition; see Supplementary Material). In general, stronger interactions (i.e., greater absolute values of *ā*) will result in greater changes in community-level sensitivity. This amplification or dampening happens because interactions modulate the rate of change in biomass with temperature across the community (altered rates of biomass accumulation), captured by the *γ* term.

In addition to this direct effect, interactions can also alter community sensitivity through the covariance term in Eq 3 (*Cov*(*w_i_*(*ā*), *E_R,i_* + *ω_i_*)), which arises when species whose biomass is strongly affected by the interactions (i.e. those with more extreme *w_i_*(*ā*) values) also tend to have higher temperature sensitivities of respiration and biomass accumulation \left(E_{R, i} +\omega_i\ right)). Although there is no empirical evidence of such relationships, it is possible that this pattern exists in nature. For example, in microbial communities it is possible that temperature and resource specialisation are positively correlated such that species with wide thermal niches (and low thermal sensitivities) also tend to be more general in their resource use (and thus are more affected by competition imposing a negative covariance structure). The relative effect of interactions through this covariance term will depend on the correlation between these two factors as well as the size of their variation across the community (i.e., greater variation in thermal sensitivity allows for more bias towards high sensitivity values).

It is important to note that our theory focuses on pairwise interactions between populations and does not explicitly consider the effects of indirect and higher-order interactions, which can be important for shaping structure and function in microbial communities^26–28^. In particular, indirect interactions in the form of intransitive (rock-paper-scissor type) competitive loops can affect coexistence and biomass dynamics in communities^27,28^. We tested the effect of intransitive interactions on our theoretical predictions and found that they had no qualitative effect on amplification of the community-level thermal response (Methods; Supplementary Fig 1). Our theoretical predictions above also do not explicitly consider higher order interactions (HOIs), where one or more non-focal species modify the direct interaction between a pair of species^26,27^. In general, we expect the effects of HOIs to alter, but not qualitatively reverse community-level amplification or dampening (Supplementary Material). Future work focusing on HOIs is needed to build a more accurate understanding of the effects of these interactions on microbial community functioning and its thermal response.

### Experimental microbial communities show an amplified thermal sensitivity of respiration rate

We tested our theoretical predictions using experiments with communities of aerobic, heterotrophic bacteria. We assembled replicated (n = 6) communities of 8 bacterial taxa (henceforth, “strains”) isolated from geothermal streams in Iceland and incubated them in a minimal, single carbon source media (M9 + glucose) at ambient temperature (20°C) using serial transfers for ~100 generations (Methods, Fig 2). This experimental design exploited the tendency of bacterial strains to increase facilitation by cross-feeding (exchanging metabolic by-products) when subject to resource limitation over time, producing replicated communities with the same strains but different interaction structures^29–31^. We henceforth refer to these as ‘adapted’ communities. As a control, we also assembled replicated communities using the same ancestral strains, but incubated them for a much shorter period of 2 days, thus limiting the time available for co-adaptation (Methods) (we refer to these as ‘*de novo* communities’). In the adapted communities, biomass dynamics stabilised after 16 days (7 transfers, ~50 generations), with 5 of the 8 strains that were originally combined persisting at a relatively stable total abundances. At the end of the experiment, after 30 days (14 transfers, ~100 generations), we re-isolated the strains from each of the communities. We assessed whether community adaptation affected the thermal response of respiration for each of the 5 strains along a broad temperature gradient (15-35°C), for populations from both the *de novo* and the adapted communities. We found no significant difference in any of the parameters of the temperature response of respiration between the ancestral isolates and the same strains isolated from the adapted communities (Fig. 3a, Supplementary Table 1), showing that the population-level temperature response of mass-specific respiration remained unchanged following adaptation, consistent with our theoretical assumption.

**Fig 2.**
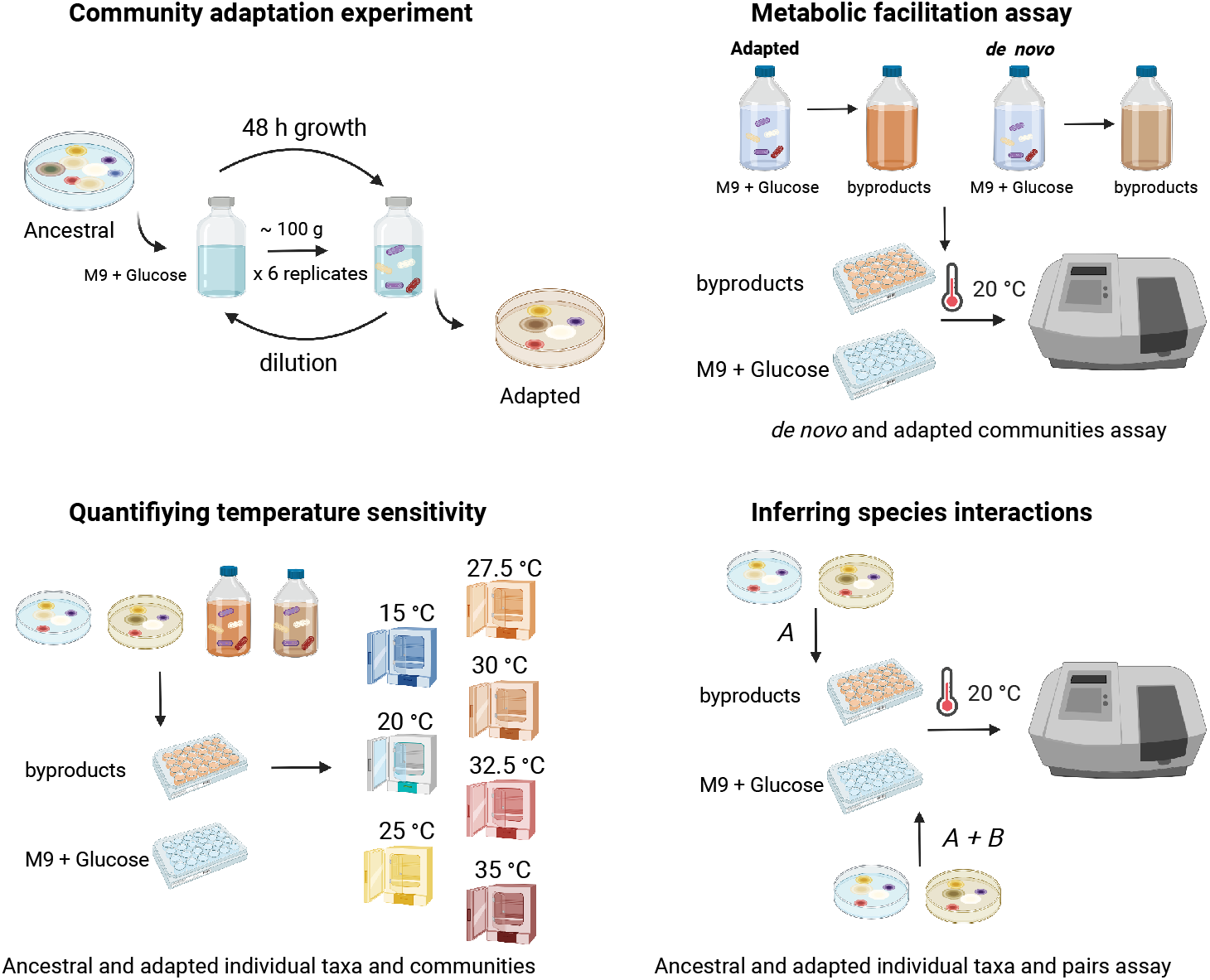
Experimental design. First, an artificial community was created using 8 bacterial strains and replicated 6 times. Communities were transferred every 48 h for 100 generations. The 5 bacterial strains coexisting after the community adaptation experiment were isolated. The 5 strains and their corresponding ancestral were incubated in M9+glucose until glucose was depleted and their byproducts preserved in sterile conditions. Ancestral and adapted taxa were incubated in M9 + glucose and the corresponding byproducts for the metabolic facilitation assay (see Methods). The ancestral and adapted individual taxa and communities were incubated in M9+glucose and the corresponding byproducts at multiple temperatures to quantify the temperature sensitivity at taxon and community levels (see methods). Finally, ancestral and adapted individual taxa and all pairwise combinations of them were incubated in M9 + glucose and byproducts for species interactions assessment. Created with BioRender.com

**Fig 3.**
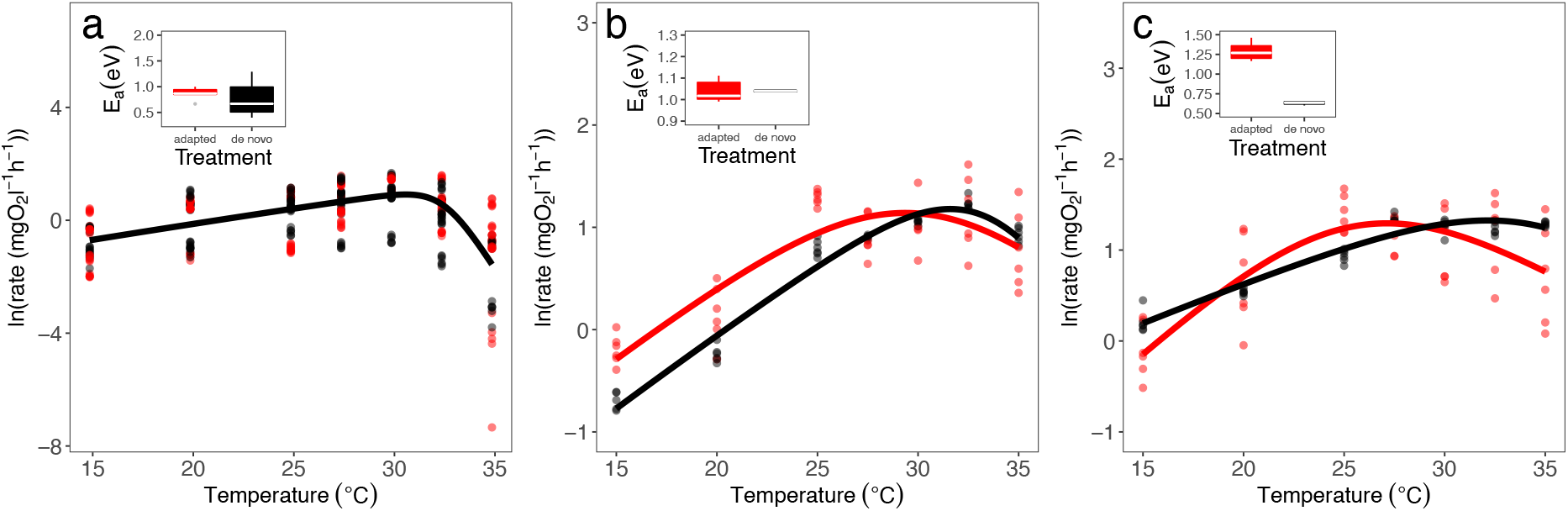
Microbial facilitation amplifies the temperature sensitivity of community respiration. Temperature dependence of respiration at the population level in M9+glucose media (a), and community level in: (b) M9+glucose and (c) spent media. Colours denote whether populations or communities are from the adapted (n= 6 biological replicates) (red) or *de novo* (n = 6 technical replicates) community isolates (black). Inset plots show the distribution of thermal sensitivities (*E*, activation energy) for each treatment where box plots depict the median (centre line) and the first and third quartiles (lower and upper bounds). Whiskers extend to 1.5 times the inter-quartile range (the distance between the first and third quartiles). The solid lines represent the average temperature sensitivity estimated by fitting the Sharpe-Schoolfield equation using non-linear mixed effects models (see Methods).

Our theory predicts that the thermal sensitivity of total community respiration should be higher in the adapted communities, where interactions were expected to have become less competitive (or more cooperative or facilitatory), compared to *de novo* communities. To test whether it was indeed facilitation through cross-feeding on metabolic by-products driving changes in interactions, we carried out community respiration assays in both M9+glucose, and “spent” media obtained by allowing communities to grow until all the initial glucose was depleted and only metabolic by-products remained (Methods). If an increase in metabolic facilitation is the main mechanism underlying changes in species interactions, then we expected to see an amplification of the thermal sensitivity of respiration in the adapted communities in the spent media, because the strains that comprise these communities would have adapted to persisting on the metabolic by-products, whereas the *de novo* assembled communities would not. By contrast, in the M9+glucose media, populations would be able to independently access glucose, relatively free from exploitative or interference competition (as the assays were carried out at low densities in the exponential phase of growth) leading to neutral interactions. Therefore, based on our theory which predicts a relatively small amplification effect when interactions are weak, for communities incubated in M9+glucose media, we expect comparable temperature sensitivities between the population- and community-level in both *de novo* and adapted treatments.

Consistent with our predictions, we found that the average temperature sensitivity of respiration was statistically indistinguishable between the *de novo* and adapted communities measured in M9+glucose (Fig. 3b, Supplementary Table 2). Furthermore, the apparent activation energy of respiration at both population- and community-levels in the M9+glucose media (where interactions are expected to be weak) were indistinguishable (species = 1.01 eV ± 0.19, community = 1.04 eV ± 0.17), as predicted by Eq 3 for near-neutral interactions. In contrast, when we quantified the temperature response of community respiration in spent media, as predicted, we found a marked and statistically significant increase in the thermal sensitivity of respiration compared to that of *de novo* communities (Fig. 3c, Supplementary Table 2), with the activation energy of the adapted community in the spent media (1.4 eV ± 0.40) 40% higher than the average population-level activation energy. To eliminate the possibility that the observed amplification of community respiration was driven by changes in per-capita, mass-specific respiration rates of the strains instead of interaction-driven biomass dynamics, we compared the thermal sensitivity of per capita respiration (mg O_2_ L^−1^ h^−1^ cell^−1^) and biomass accumulation (methods) between the *de novo* and adapted communities. We found that the thermal sensitivity of per-capita respiration was indeed statistically indistinguishable between the *de novo* and adapted communities (Fig. 4a-b, Supplementary Table 3), while that of biomass accumulation was significantly higher in the latter when communities were incubated in spent media (Fig. 4c-d Supplementary Table 4).

**Fig 4.**
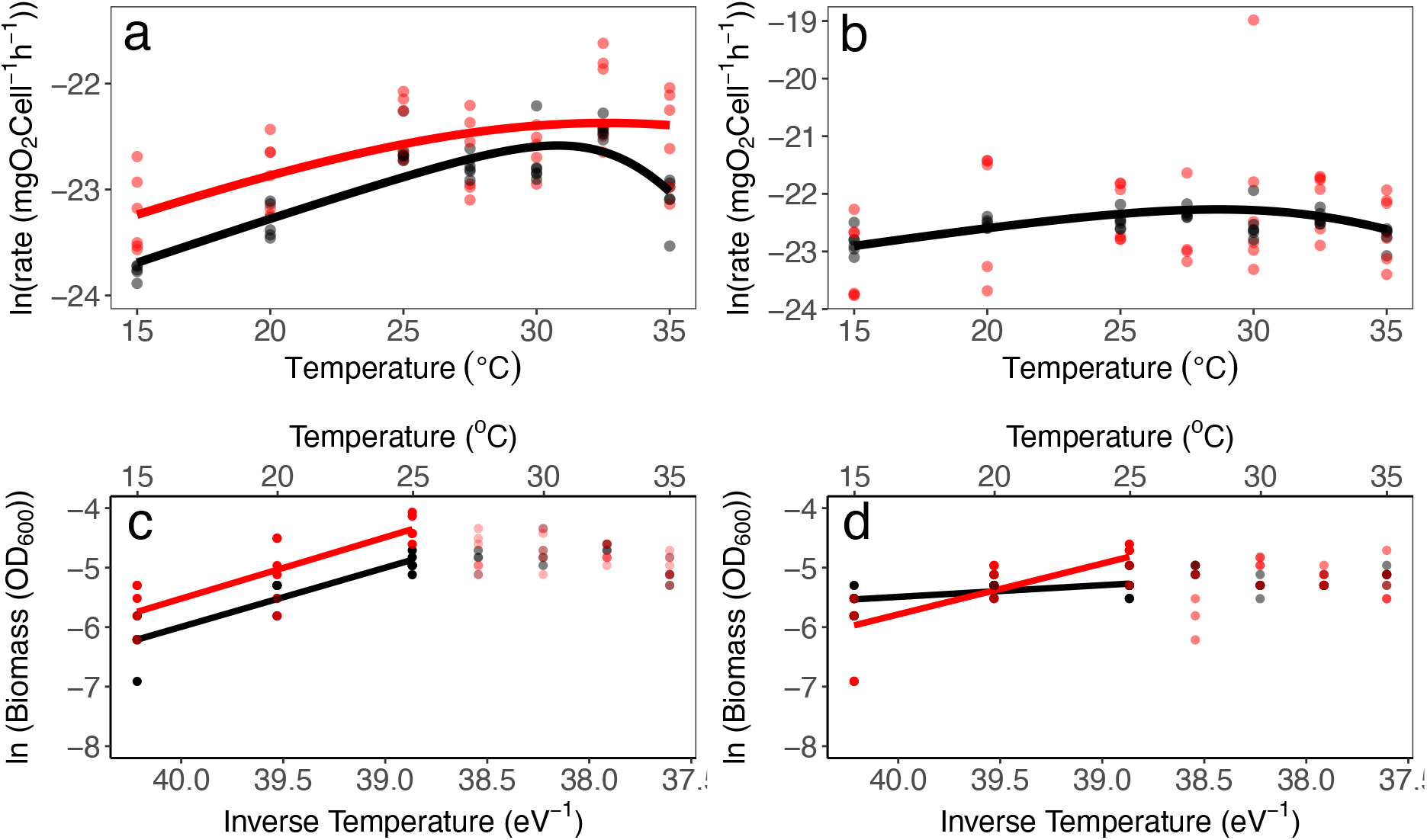
Temperature dependence of respiration per cell and community biomass. Upper panels show respiration per capita (ln respiration rate (mg O2 cell^−1^ h^−1^)) at different assay temperatures (15, 20, 25, 27.5, 30, 32.5 and 35 °C) for *de novo* assembled (black) and adapted (red) communities in M9 + glucose media (panel a) and ‘spent media’ (SM) (panel b). Details about model fitting and parameters estimation are given in Supplementary Table 3. Lower panels show community biomass (ln (OD_600_)) at different assay temperatures (15, 20, 25, 27.5, 30, 32.5 and 35 °C) for the *de novo* assembled (black) and adapted (red) communities in M9 + glucose media (panel c) and ‘spent media’ (SM) (panel d). Bright coloured points represent the exponential part of the temperature dependence and transparent points represent the biomass estimates beyond the maximum which were not included in the estimate of the temperature sensitivity. Details about model fitting and parameter estimation are given in Supplementary Table 4.

### The amplification in thermal sensitivity is driven by a shift from competition to facilitation during community adaptation

Next we confirmed that interactions had indeed become more facilitatory (or less competitive) in the adapted relative to the *de novo* communities through two additional experiments. First, we first looked at how the asymptotic biomasses of the communities and the individual strains (i.e., the carrying capacity, *K*) changed following adaptation. If interactions between populations had become more facilitatory, we expected to see higher biomass attained in these communities due to more efficient use of limiting resources. When each taxon was grown in monoculture on the M9+glucose media for 72 h at 20°C, we saw no statistically significant effect of treatment on *K* before and after adaptation (p=0.56, Supplementary Fig 2, Table 5). However, when the same test was performed with communities, *K* was higher in the adapted communities (Fig. 5a, M9+glucose media, Supplementary Table 6). To further investigate these effects, we repeated the experiment, but this time incubated the communities in spent media. We found that the adapted communities reached higher *K* compared to the *de novo* assembled communities in the spent media (Fig 5a, Supplementary Table 6). Furthermore, the *K* attained by the adapted communities in the spent media was even higher than they attained in the M9+glucose media. Thus, co-adaptation between strains increased the biomass production efficiency of the community for the given set of resources. Because we observed an increase in community-level performance of the adapted communities in M9+glucose media and spent media, it is likely that the facilitatory (or less-competitive) interactions persist in M9+glucose, albeit with a smaller impact on community biomass because on average, each strain’s population relies less on metabolic by-products in resource-rich media.

**Fig 5.**
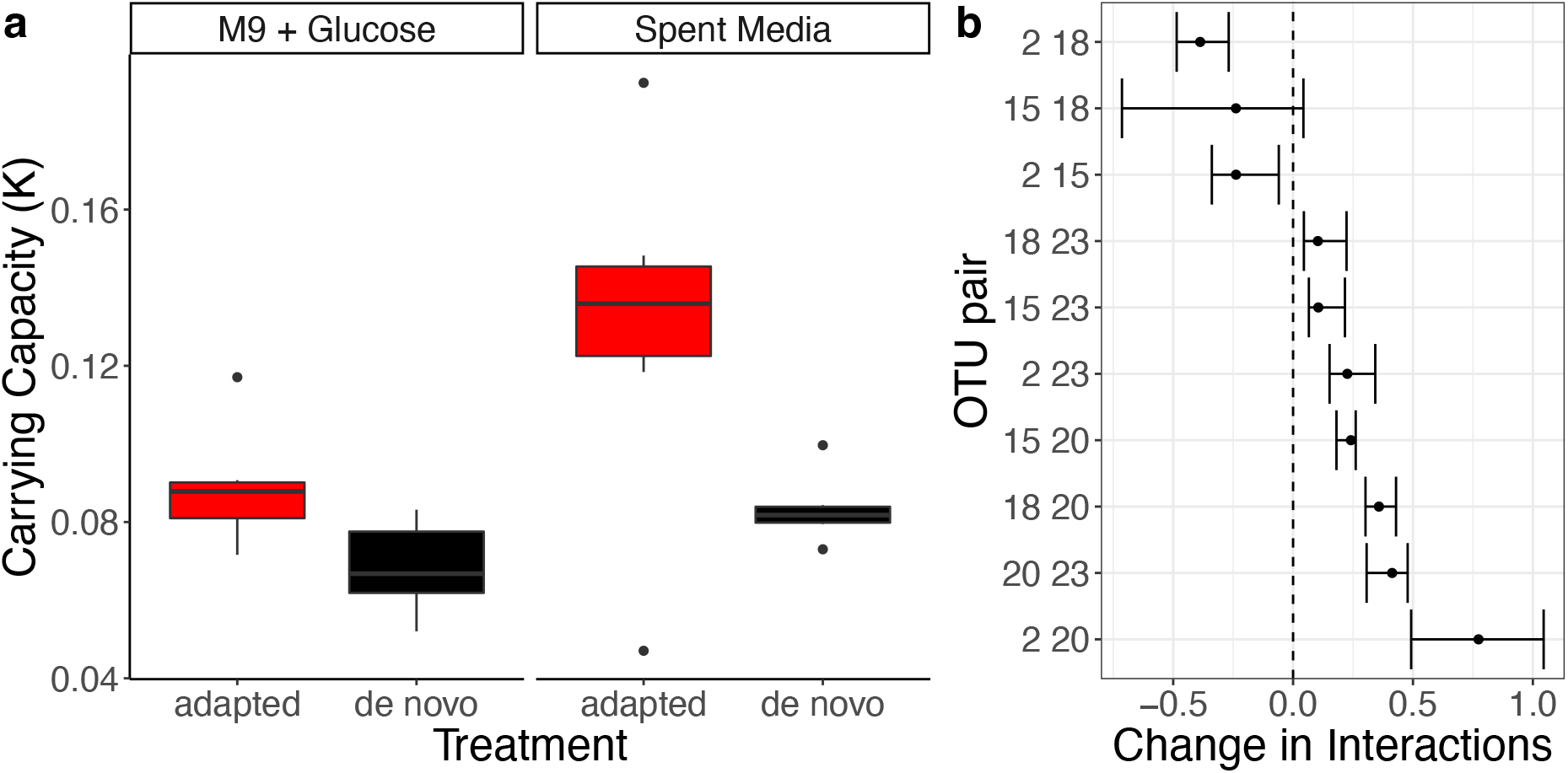
Emergence of facilitation in assembled synthetic microbial communities. (a) Carrying capacity of the adapted (n = 6 biological replicates for each media treatment) and *de novo* (n= 6 technical replicates for each media treatment) assembled communities grown in either M9+glucose or spent media for 72 h at 20°C (Supplementary Table 6). Box plots depict the median (centre line) and the first and third quartiles (lower and upper bounds). Whiskers extend to 1.5 times the inter-quartile range (the distance between the first and third quartiles). (b) Change in interaction strength between pairwise combinations of strains from the *de novo* to adapted communities in the M9+glucose. Each row shows the average and 95% bootstrapped confidence intervals of the estimated change in interactions for each pair. Negative values indicate interactions have become more competitive while positive values indicate that interactions have become more facilitatory. The interactions become predominantly more facilitatory (70% of the cases), with *<u>a</u>* changing from −0.07 in the *de novo* to 0.065 in the adapted communities.

We then experimentally quantified the change in direction and magnitude of pairwise species interactions after adaptation. We grew isolates of each strain individually, and in all possible pairwise combinations of strains, before and after adaptation for up to 72 h (depending upon when they reached carrying capacity) at 20°C in M9+glucose and the spent media. We then estimated the mean of the pairwise interaction strengths using a measure based on the comparison of growth rates in mono-versus paired-cultures (Methods; Supplementary Figure 3). When grown in M9+glucose media, interactions predominantly shifted towards positive values following adaptation indicating a shift from competition to facilitation (Fig. 5b). When the same experiment was attempted in mono-culture in spent media, many of the isolates did not grow despite the whole communities being able to coexist under these conditions. This indicates that the resource environment that emerges from the metabolic by-products in multi-species communities is essential for the co-adapted populations to grow. The metabolic by-products of just a single community member is insufficient to enable persistence. This point can be made more exact by analysing the Lotka-Volterra equations (Methods). Our experimental results showed that the population growth of monocultures in spent media was negative, such that growth, plus the effects of density dependence, were negative, i.e., *r_i_* − *a_ii_C_i_* < 0. The joint population growth is still negative in paired-cultures because the effect of positive pairwise interactions is still not enough to allow persistence. In contrast, the whole community’s growth is positive where all interspecies interactions are present, i.e., *r_i_* − *a_ii_C_i_* + *N<u>a</u> <u>C</u>* > 0. Taking both these constraints into account, it follows that the net effect of interactions must be positive in order to outweigh the negative intrinsic growth rates, and that *<u>a</u>* > 0 in the assembled communities, consistent with our estimates of the pairwise interaction coefficients. We note that it was not possible to estimate each of the pair of (potentially asymmetric) interaction coefficients between any two strains in paired-culture in our experimental system, because this would have required us to track the relative abundance of each taxon over time. Importantly, our prediction that a shift towards positive species interactions amplifies the thermal sensitivity of community respiration pertains to the overall average of interaction strengths in the community, for which an estimate of the mean interaction strength between pairs of strains is sufficient.

Finally, we tested the possibility that a change in the interaction structure to favour populations with higher thermal sensitivity (meaning an increase in the covariance term *Cov*(*w_i_*(*<u>a</u>*), *E_R,i_* + *ω_i_*) in Eq. 3) was responsible for the amplification of community respiration by examining the relationship between the population-level thermal sensitivity values and estimated interaction coefficients. We found no relationship between these two features (Supplementary Figure 4), either before or after adaptation, indicating that the covariance term contributed relatively little to the overall temperature sensitivity of community respiration. That is, the increase in thermal sensitivity of community respiration cannot be explained by an increase in the tendency for interactions to positively affect populations with high thermal sensitivity.

## DISCUSSION

In summary, our theoretical and empirical results provide compelling evidence that a shift from competition to facilitation results in positive population dynamical feedbacks that ultimately amplify the thermal sensitivity of community-level respiration rate by increasing total biomass. This amplification occurs in a predictable way, and can be quantified through a general relationship between the magnitude and direction of average interaction strength in the community and the thermal sensitivity of its respiration that we have derived here.

Our finding that changes in the direction and strength of microbial species interactions can so profoundly alter the temperature sensitivity of community-level respiration, has far-reaching implications given the significant contributions that microbial communities make to ecosystem functioning in aquatic and terrestrial environments. Microbial communities tend to have either competitive or cooperative interactions across space^20^, or over time (during community assembly for example^30,32^). This implies that microbial communities can either dampen or amplify the effects of temperature change on carbon cycling. For example, changes in interaction structure over time due to longer-term assembly and turnover dynamics could mean that the same microbial community would switch between states that dampen (when competition dominates, at early stages of community assembly) and amplify (with facilitation dominates, in later states of community assembly) the sensitivity of ecosystem functioning to temperature change. In particular, microbe-mediated decomposition of organic matter, which is the main contributor to CO_2_ and CH_4_ fluxes in the carbon cycle, depends on facilitation among taxonomically diverse consortia of bacteria and archaea. Indeed, our finding that weakening of competition and strengthening of facilitation amplifies the temperature sensitivity of community metabolism may help to explain the relatively high thermal sensitivity seen in syntrophic methanogenic microbial communities^33^.

## METHODS

### Deriving the thermal sensitivity of ecosystem respiration

For a given ecosystem (henceforth, synonymous with a bacterial community), total temperature-dependent respiratory carbon flux, *R_eco_*(*T*) can be expressed as the sum of the products of species (strain-)-level mass-specific respiration rates *R_i_*(*T*)’s and biomasses *C_i_*(*T*)’s (Eq (1)). We are interested in the thermal sensitivity of ecosystem respiration, *E_eco_*:

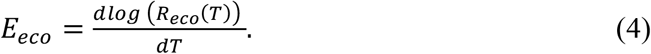

Using the fact that 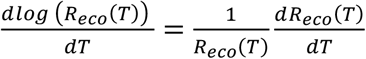 and substituting Eq (1) in Eq (4), we get:

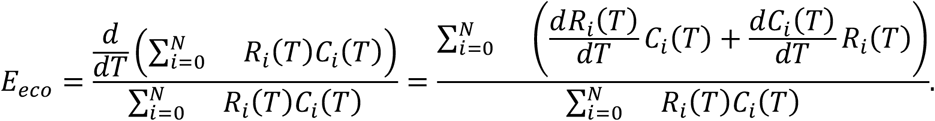

Using the general definition 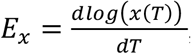, this simplifies to Eq 2. Equation 2 shows that the thermal sensitivity of ecosystem respiration is given by the average of the sensitivities of biomass and respiration to temperature across the community, weighted by the respiratory output of each species. We can further explore this by defining 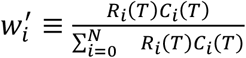 (effectively, a normalised weighting parameter) which lets us write *E_eco_* as:

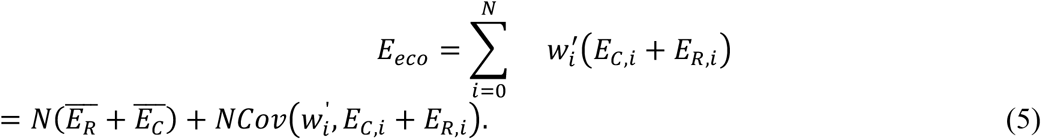

where *E_C_* and *E_R_* are the average thermal sensitivities of biomass and respiration across all *N* species in the community (defined as 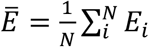). Expressing *E_eco_* in this way shows that it depends on both, the average sensitivities across the community and the covariance between the weightings (i.e. the relative contribution of each species to total respiration) and the thermal sensitivities of individual species. Note that this partitioning of total ecosystem function into the contributions of the average effect across populations and the specific structure of their contributions and biomass (the covariance term) is the same as use in the Price equation which has been previously applied to understand ecosystem function and the effects of species loss^34^.

Next, we consider how inter-species interactions affect the thermal sensitivity of ecosystem respiration. Because species interactions are expected to change species’ biomasses much more rapidly than respiration rates (which change at much longer, macro-evolutionary timescales^35^), we focus on the effects of interactions on the biomass components of Eq 2 (the *C_i_*’s and *E_C,i_*’s). To model these effects, we use the generalised Lotka-Volterra (GLV) model:

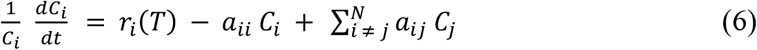

Here, *r_i_*(*T*) is the (temperature dependent) intrinsic growth rate of the *i*th species, *a_ij_* the strength of the effect of the *j*th species on the *i*th one (positive or negative), and *a_ii_* the strength of (negative) intraspecific density dependence. For an arbitrary structure of species interactions (signs and strengths of the *a_ij_*’s), it is impossible to meaningfully determine how interaction structure affects species’ biomasses. Therefore, next, we derive an approximate relationship with the aim to quantitatively predict the response of biomass in the early stages of community assembly as follows.

First, we derive a mean-field approximation^36,37^ of the GLV model (Eq 6). We use the definition of the average interaction experienced by a focal species, 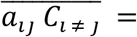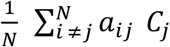 to write the interspecies interactions term in Eq 6 as:

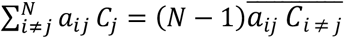

Assuming the system is large (*N* is large) and the difference between 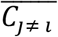 and 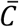 (i.e. the: exclusion of the *i*th species has little effect on 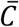), we express the interaction term as:

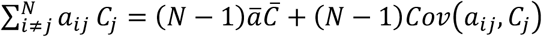

Assuming that the effect of any single interaction on a species’ biomass is small, we can take the covariance term to be negligible, yielding the mean-field approximation of the GLV (Eq 6) model:

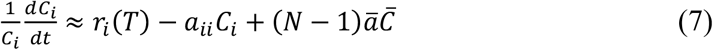

Although this approximation relies on the assumption of large community size, we show that our results hold qualitatively even in small communities (Supplementary Figure 5).

Next, we need to solve this system of equations representing an ecosystem for the *C_i_*’s, in order to determine the (temperature-dependent) effects of species interaction structure on population biomasses, *C_i_*. We use a Taylor-series expansion around *t* = 0 to get an approximate expression for *C_i_* in the early stage of community assembly in terms of average interaction strength *a*. We start by considering log-biomass as a function of time, log(*C_i_*(*t*)), which can be approximated around *t* = 0 giving:

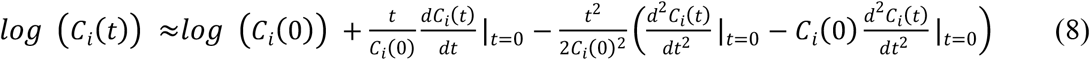

We use Eq 8 to get the second order derivatives (by taking the time derivative again). This in turn requires an expression for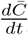, which we obtain using the Taylor-series approximation of the average of a function of uncorrelated random variables *x_i_,…,x_N_*,

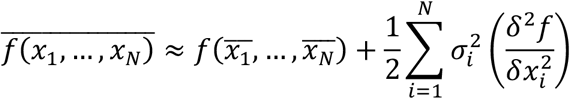

Combined, these give an expression for the *i*th species’ log biomass at time *t*:

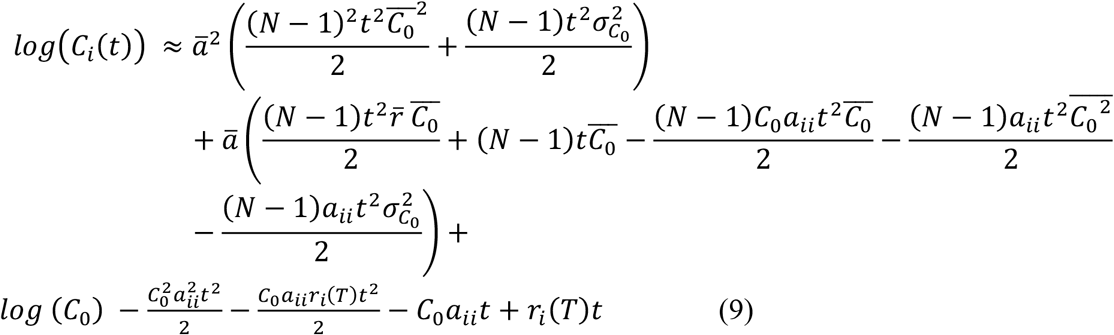

We can take the derivative of Eq 9 with respect to temperature to get an expression for *E_C,i_*:

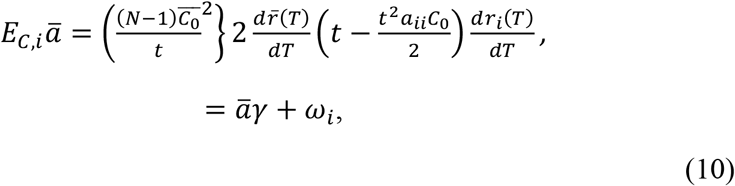

Where 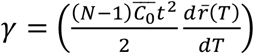 and 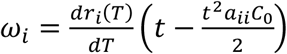 are constants representing the temperature dependence of average biomass growth across the whole system and the biomass growth of species *i* respectively.

Finally, substituting Eqs 10 back into 5 gives an expression for the thermal sensitivity of respiration across the whole system (same as Eq 3):

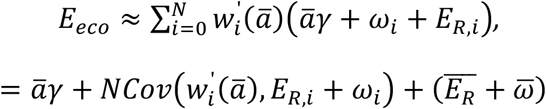

Supplementary Figure 6 shows that this approximation adequately captures the qualitative effects of interactions on thermal sensitivity of ecosystem respiration. Thus, as explained in the main text, the above equation predicts that *E_eco_* will depend only on internal factors affecting species’ population growth (growth rates and intra-specific interactions) when *ā* = 0 (competitive and facilitatory interactions balance each other), and relative to this, it will be dampened if *ā* < 0 (competitive interactions dominate) and amplified if *ā* > 0 (facilitatory interactions dominate).

#### Generating specific predictions

To generate specific predictions based on this theory, we parameterised Eq 9 with randomly generated communities of *N* = 50 to obtain biomass estimates in the early stages of assembly (*t* = 3.0), setting *C*(0) = 0.01. We consider this time frame as it corresponds to the experimental setup. For each such synthetic community we multiplied these biomass estimates by mass-specific respiration *R* and summed over all populations to get the total ecosystem respiration *R_eco_* (Fig 1b). This process was repeated at different fixed temperatures, from 5–25°C. We used a modified Boltzmann-Arrhenius equation to represent the temperature-dependence of growth *r* and respiration *R* rates,

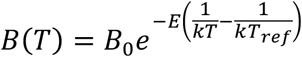

where *B*_0_ is a normalisation constant, *E* is the temperature sensitivity, *K* is the Boltzmann constant, and *T* and *T_ref_* are the temperature and reference temperature (set to 15°C) respectively. For both *r* and *R*, we sampled the *B*_0_ and *E* values from normal distributions such that *B*_0_ ~N(1,0.1) and *E*~N(0.6,0.1). We considered three types of community interaction structures—competitive, neutral and facilitatory— by setting *a* = −0.02, 0, and 0.02 respectively. Intraspecific interactions were all set to *a_ii_* = −10. We calculated *E_eco_* by using the same parameters in Eq 3 to generate the values in the inset plot of Fig 1b.

### Isolation and identification of bacterial taxa

The experiment was conducted with 8 bacterial taxa isolated from a geothermal valley in Iceland (see Supplementary Table 7). These were isolated from biofilm samples collected from the surface of rocks in May 2016-May 2017 in Hvergerdi Valley, 45 km east of Reykjavik, Iceland. Samples were immediately frozen upon collection with 17% glycerol and transported at −20°C for further processing in the laboratory. Upon return to the laboratory, samples were thawed at 20°C and prepared by serial dilution and plating 10 μL onto R2A agar plates (Oxoid Ltd) with sterile glass beads. Plates were incubated at a range of temperatures between 15-25°C for 5-10 days. The resulting colonies were distinguished by morphology, picked and placed into 200 μL Lysogeny Broth (LB) and incubated for 48 hours. To preserve the library of taxa, samples were then centrifuged, the supernatant was removed and the pellet was re-suspended in mix of LB and 17% glycerol before being frozen at −80°C.

Isolates were assigned taxonomy using 16S Polymerase Chain Reaction (PCR) followed by Sanger sequencing within the 16SrRNA gene. A master-mix solution was prepared using 7.2 μL of DNA free water, 0.4 μL 27 forward primer, 0.4 μL 1492 reverse primer and 10 μL of Taq polymerase per sample. To create a template solution 2 μL of sample 100x diluted in DNA free water was added to 18 μL of master-mix solution. Samples were then placed in a thermal cycler (Applied Biosystems Veriti Thermal Cycler). This procedure included 1 cycle at 94°C for 4 minutes, 35 cycles at 94, 48 and 72°C for 1 minute, 30s and 2 minutes, respectively, and finally, 1 cycle at 72°C for 8 minutes. The PCR product was cleaned up using Exonuclease I and Antartic Phosphatase and high-quality samples were Sanger sequenced using the 27F, 1492R primers (Core Genomic Facility, University of Sheffield). Sequences were trimmed in Genious (version 6.1.8) removing the bp from the 5’ end and trimming the 3’ end to a maximum length of 1000bp. Using Mothur v.1.39.5^38^, sequences longer than 974bp were aligned to the Silva.Bacteria. Taxonomic identities were assigned using the RDP trainset 9 032012 as a reference database (Supplementary Table 7). Morphology was assessed visually to allow recognition of each taxon when mixed in an experimental community.

### Community adaptation experiment

We assembled replicated communities with the 8 bacteria taxa. Stock cultures were first grown in LB medium at 20°C overnight to establish a dense, healthy culture and then standardized to a common biomass density in M9 media with 0.2% glucose. We used this minimal growth medium because it has a single, defined, and easily quantifiable carbon source. A “community stock” solution was built by adding 100 μL of each taxon. Then, an 40 μL aliquot of the community stock solution was added to 5000 μL of M9 media + 0.2% glucose and incubated at 20°C in glass vials for 48 h. We used 6 replicates and 2 blanks to check for media contamination. We transferred each community every 48 h by diluting 40 μL of the community in 5000 μL of fresh media. Each transfer encompassed both exponential and stationary (typically reached within 24h) phases of the growth cycle. Preliminary experiments revealed that over 95% of the glucose was depleted after 24h. Consequently, the duration of each transfer encompassed both resource replete conditions where glucose was abundant, and periods where glucose was scarce. In the resource scarce periods, persistence of strains was expected to be strongly dependent on their ability to utilise recycled carbon in the form of metabolic by-products (cross-feeding). Each community was passaged 14 times (30 days, ~100 generations) in this manner over the course of the experiment. Optical density (OD_600_) of the community was measured at every transfer using a Themo Scientific™ Multiskan Sky Microplate Spectrophotometer at 600 nm. Each community was also plated on R2A agar once a week to ensure they remained uncontaminated. At the end of the experiment, each community was plated and individual taxa isolated, identified and stored at −80°C for downstream analysis. We henceforth refer to these replicate communities, established from a common pool of taxa and then incubated across multiple generations under intermittent resource-depleted conditions, as “adapted”. For comparison, we assembled six replicated communities with the same 8 taxa in the same conditions as the adapted communities without transferring (passaging) them several times. We refer to these communities as “de novo”.

### Metabolic facilitation assay

To investigate whether metabolic facilitation emerged in each community we carried out an assay to quantify levels of total biomass production in conditions where the only available resources were metabolic by-products generated during its assembly (“spent media”). We hypothesised that if the development of metabolic facilitation were to drive species interactions to be more positive in communities where the constituent members had been grown together under resource limitation, then we should observe enhanced biomass production in the adapted compared to de novo communities, when both were grown in spent media. This is because metabolic facilitation should enable better growth on the metabolic by-products of community members. For this, the de novo and adapted communities were each inoculated into 5 mL of M9 media + 0.2% glucose and incubated at 20°C until the glucose level in the media fell below detection limit (typically, 48 h).

A Glucose GO Assay Kit (Sigma) was used to monitor the level of glucose in the media. The samples were then centrifuged at1421 g (3000 rpm), for 10 min. We refer to ‘spent media’ as the resulting media containing species metabolites without any other carbon source. The spent media was filter-sterilized and stored at 4°C. The de novo communities were then re-grown in the spent media extracted from their own (shorter) incubation. Each unique adapted community was re-grown in its own spent media. Biomass was measured every 4 hours as optical density using a Themo Scientific™ Multiskan Sky Microplate Spectrophotometer at 600 nm.

The community-level carrying capacity was quantified by fitting the logistic growth model to the resulting time-series using non-linear least squares regression using the R package `nlsLoop` (following Garcia *et al.*^39^). At the single-taxon level, we used an analysis of variance (ANOVA) to test for significant differences in carrying capacity among treatments (‘ancestral’ or ‘adapted’) (Supplementary Table 5). At the community level, we also used an ANOVA to test whether carrying capacity differed significantly between treatments (‘adapted’ or ‘de novo’), media sources (‘M9+glucose’ or spent media) or their interaction (Supplementary Table 6).

### Quantifying the temperature sensitivity of respiration

We characterized the thermal response curves for respiration at the taxon level for both, the ancestral and adapted strains (isolated from the adapted communities). The isolates were grown overnight in LB medium from −80°C freezer stocks, then transferred into M9 media + 0.2% glucose and acclimated in incubators at 9 temperatures (15°C, 20°C, 25°C, 27.5°C, 30°C, 32.5°C, 35°C, 40°C, 45°C) for 24 hours. The incubation time was selected based on the time the isolates take to reach carrying capacity. After acclimation, biomass was estimated by measuring optical density using a Themo Scientific™ Multiskan Sky Microplate Spectrophotometer at 600 nm and then standardized at the same optical density (OD_600_ = 0.05). A 4 mL aliquot of each sample was added into 46 mL of fresh M9 media. We used 6 technical replicates of each taxon for each treatment (de novo & adapted) and took the average as our estimate of respiration rate. All measurements were made while the isolates were in the exponential phase of growth.

At the community level, we measured the temperature sensitivity of respiration for the adapted and de novo assembled communities. The de novo isolates were grown in LB medium overnight immediately after coming out of the −80°C freezer, then transferred into either M9 media with 0.2% glucose or the spent media containing only metabolic by-products. Each of the 6 adapted communities (i.e. 6 biological replicates) were transferred directly to fresh M9 media with 0.2% glucose or the corresponding spent media. All samples were acclimated in the appropriate assay media (either M9+glucose or spent media) in Percival incubators at 9 temperatures (15°C, 20°C, 25°C, 27.5°C, 30°C, 32.5°C, 35°C) for 24 hours. After acclimation samples were standardized to the same biomass (OD_600_ = 0.05). The de novo communities were then assembled by adding 800 μL of each ancestral isolate into 46 mL of the corresponding media (M9+glucose or spent media). Each de novo community was replicated 6 times. For the adapted communities, we added 522 μL of each replicate community into 6 mL of the corresponding media (M9+glucose or spent media).

#### Respiration rate measurements

Respiration was measured as oxygen consumption using an array of 10 SensorDish Readers (SDR, PreSens GmbH, Regensburg, Germany). Each plate reader can analyse 24 samples, meaning the array of 10 allowed us to measure 240 samples simultaneously. The PreSens system was calibrated using a two-point calibration at each measurement temperature. 0% oxygen saturation was defined using a solution of 1% (w/w) sodium sulfite and 100% oxygen saturation used air-saturated water. In each well we placed a 5 mL vial with the sample to be measured. The vials were slightly overfilled so that no air was trapped within the vials as the lids were closed. The equipment was then run in parallel at the 9 temperatures (see above) and measured the concentration of dissolved oxygen every minute for ~4 h. The rate of respiration was derived from the slope of a linear regression of oxygen concentration against time (mg O_2_ l^−1^ h^−1^). To estimate respiration per cell, a 200 μL aliquot for each treatment, media and replicate was sampled after measuring the respiration rate to quantify bacterial abundance. Samples were fixed with paraformaldehyde and glutaraldehyde (P+G) 1% final concentration and kept in the −80 °C freezer. Samples were stained using Sybr™ gold nucleic acid stain and analysed using the BD Accuri™ C6 flow cytometer in low flow rate. Total community respiration (mg O_2_ l^−1^ h^−1^) was divided by the total bacterial abundance (cell l^−1^) in order to obtain the respiration per cell.

#### Biomass estimates

To estimate community biomass, a 200 μL aliquot for each temperature, treatment, media and replicate was sampled after community respiration rate measurements were completed. The samples were then analysed in a Themo ScientificTM Multiskan Sky Microplate Spectrophotometer standard plate reader at 600 nm to obtain the total bacterial biomass in OD_600_. A blank containing the media without any bacterial cell was also analysed to correct the data by subtracting the blank to the sample value.

#### Model Fitting

We fitted the four parameter Sharpe-Schoolfield equation to the respiration rate data measured along the thermal gradient to calculate thermal sensitivity (see Padfield *et al*.^40^). At taxon level, we fitted the Sharpe-Schoolfield model to the rate data using nonlinear mixed effects models in the `nlme` package in R. We modelled species as a random effect and treatment (ancestor or adapted) as a fixed effect on each parameter. Model selection started with the most complex possible model, including fixed effects on all parameters and then proceeded by removing treatment effects on each of the parameters. At the community level, we fit the Sharpe-Schoofield model separately to each media (M9+glucose and spent media) using nonlinear mixed effects models in the `nlme` R package. We modelled replicate as a random effect and treatment (de novo or adapted) as a fixed effect on each parameter in the Sharpe-Schoolfield equation. With both analyses, model selection started with the most complex possible model, including fixed effects on all parameters and then proceeded by removing treatment effects on each of the parameters. We used likelihood ratio tests for model comparisons (see Supplementary Tables 1-3).

The temperature dependence of total biomass was quantified using the Arrhenius equation by applying a linear mixed effects model to the natural logarithm of total biomass along the exponential part of the thermal response (15 to 25°C). We used only the exponential part of the thermal response curve as total biomass did not follow a typical unimodal shape which precluded fitting the Sharpe-Schoolfield equation.

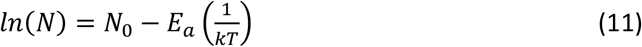

Where N is community biomass (ln (OD_600_), N_0_ is the rate constant, Ea is the activation energy, k is Boltzmann’s constant (8.62×10−5 eV K^−1^) and T is the absolute temperature in °C. We modelled replicate as a random effect on the intercept and treatment (de novo or adapted) as a fixed effect on both the intercept and the slope (Ea), estimates are given in Supplementary Table 4.

### Inferring species interactions

To quantify how long term co-culture altered the nature and strength of biotic interactions we use an approach based on the difference in experimentally observed growth rates when strains are grown individually versus when they are grown in pairs. In brief, our approach involves two stages. First, using the microcosm experiments we estimate growth rate for each strain individually, and in all pairwise combinations. We then use these to estimate the direction and magnitude of interactions based on a model of population growth detailed below.

To measure growth rates for both pairs and individual strains, we grew isolates of each taxon before and after adaptation at 20°C in M9+glucose in monoculture, as well as in all possible pairwise combinations of taxa. For each incubation, we measured OD_600_ every hour until each pair reached carrying capacity. Depending on the species and treatment this incubation time could vary between 24 to 72 h. We then repeated the experiment using the spent media (see “Metabolic facilitation Assay’’ above). This second set of experiments was focused on growing each taxon in the metabolites of the others in all pairwise combinations. First, we grew the five original isolates in LB medium. All taxon abundances were standardized at OD_600_ = 0.1 diluted into M9+glucose. We then inoculated 40 μL of each taxon’s population into 5 mL of M9 media with 0.2% glucose, in 5 mL vials. We incubated each vial until there was no detectable glucose remaining. The cells were separated from the spent media by centrifuging in 15 mL falcon tubes at 3000 rpm for 10 min and the spent media was filter-sterilized and stored at 4°C. We used the same protocol with the adapted isolates to obtain a spent media with their metabolic byproducts. Subsequent experiments with the ancestral or adapted isolates used the corresponding spent media. We inoculated each of the five taxa in M9+glucose media and in the spent media of the five taxa in all pairwise combinations at 10% v/v in a 384 well plate. All taxa were standardized at OD_600_ = 0.05 before being inoculated to 90 μL of M9+glucose or spent media. The plate was incubated in a Themo Scientific™ Multiskan Sky Microplate Spectrophotometer plate reader at 20 °C and OD_600_ was measured every hour until carrying capacity was reached. To obtain estimates of growth rates (*r, h^−1^*) from both experiments we fit both logistic and exponential growth models to the OD_600_ data from the first 20 hours. This time limit was chosen because it encompassed the growth phase (i.e. either before populations reached carrying capacity, or started to decline) across all strains and pairs. The growth estimate was taken from the best fitting model as indicated by the lowest AIC for each strain or pair and treatment combination.

We used the growth rates obtained from these experiments to estimate pairwise interaction coefficients using a method based on the difference in growth rates when strains are grown in pairs or in monoculture. This method is similar to that previously used to assess the nature of interactions in bacterial communities^29,39^ (see Supplementary Materials for further discussion) and is derived as follows.

First, consider the growth of a pair of strains *x*_1_ and *x*_2_ in isolation, which can be modelled by the growth equations:

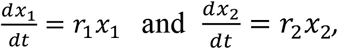

where *r*_1_,*r*_2_ are the mass-specific growth rate and *x*_1_,*x*_2_ are the biomass of each population. Note that we omit the intraspecific density dependence terms here as we are interested in the early stages of population growth within which the experimental observations were made. At these timescales the effects of density dependence will be of order O(*C*^2^), and thus smaller than the effects of actual growth rates, allowing them to be left out of the growth equation. In practice, any deviations from the true maximal growth rate caused by density dependence in the experiments will be captured in the effective growth rate we measure.

When both strains are grown together the additional interaction terms *a*_12_ and *a*_21_ need to be introduced to capture the effect of interactions between the two species:

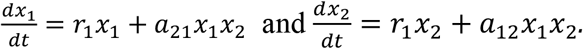

Next, we write the equations for the growth the total biomass of the pair *x_tot_* as:

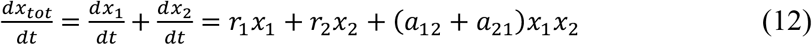

which we can also approximate in the early stages of the pair’s assembly using a single effective growth rate which incorporates the effects of both intrinsic population growth and interactions from Eq 12. This is done in terms of the total biomass of the pair *x_tot_* which the only quantity observable in the experimental data:

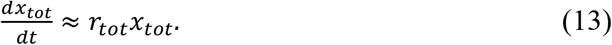

Combining Eqs 12 and 13 thus gives:

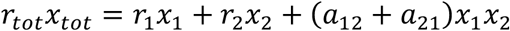

allowing us to solve for the total interaction strength:

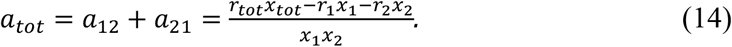

Equation 14 defines a line of solutions on which the values of *a*_12_ and *a*_21_ can lie.

Taking into account that at the beginning of the experiment each strain is at equal abundance i.e.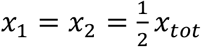 and 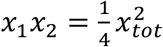 we can write the total interaction strength as:

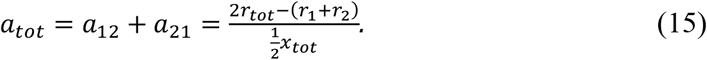

Equation 15 shows how the total interaction strength is given by the deviation of the total paired growth from the null-case with no interactions (given by the sum of the individual strain growth rates in monoculture). If the pair grows at a lower rate than expected from their growth in monoculture then we infer that the interaction between them is competitive and vice versa. Note that the use of the biomasses at *t* = 0 here is not to say that biomasses of strains are constant over time but rather that the relative contributions of the growth rates of the individual strains to the null, non-interacting case is equal. In this way the biomasses are only used to correctly weigh the individual monoculture growth rates in the null model. The change in biomass over time is captured in the growth rate terms and we do not assume that the biomasses are constant.

In order to derive an estimate of the individual pairwise interaction coefficients we next consider the case where interactions are symmetric *a*_12_ = *a*_21_ = *α* letting us write:

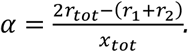

Note that this assumption is the same as considering the average interaction strength of the pair: 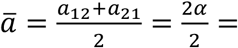. Therefore, as such, the symmetry of interaction coefficients does not affect our inference of the *overall average interaction strength across the community*. In order to calculate asymmetric interactions, one would require data on the relative abundances of strains when grown together, over time.

As the total abundance is held constant across all the experiments, the *x_tot_* term in the denominator acts as a single scaling term across all interaction estimates and can thus be dropped giving the final expression:

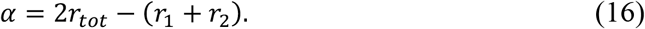

To apply Eq 16 to our data, we use a bootstrapping procedure to account for variation in growth rate estimates amongst replicates. Specifically, taking the data from the growth curve experiments, we sample each of the parameters from Eq 16 with replacement across the replicates 10,000 times and calculate *α* for each taxon pair. This gives a distribution of estimates of *α* for each pair as shown in Fig 5.

## Data and Code Availability

All data and code to reproduce our results are at https://doi.org/10.5281/zenodo.7105128

## Acknowledgements

This work was supported by a European Research Council Starting Grant awarded to G.Y-D (ERC StG 677278 TEMPDEP). T.C was supported by the QMEE CDT, funded by NERC grant number NE/P012345/1. S.P was funded by Leverhulme Fellowship RF-2020-653\2 and UK national NERC Grants NE/M020843/1 and NE/S000348/1.

## Author Contributions

G.Y-D and S.P. conceived the study; F.G. and G.Y-D designed the lab experiments; F.G., R.W. and D.B.O carried out the lab experiments, T.C and S.P developed the theory; all authors conducted the analysis of the experimental data and wrote the manuscript.

## Additional Information

