## Supplementary Materials for "A shift from competition to facilitation amplifies the temperature-dependence of microbial community respiration"

**Potential effects of higher-order interactions.** The GLV equations (Eq 6) are in fact an "effective" system obtained by approximating the dynamics of the more complex underlying consumer-resource system, which allows the interactions between populations through resource dynamics to be represented by direct, pairwise coefficients[41,42]. These interaction coefficients thus embody the combined effects arising through resource uptake and cross-feeding on metabolic by-products. Letten & Stouffer [30] have shown that in cases where resource dynamics are not captured by the GLV approximation, its accuracy can be improved by including higher-order interaction (HOI) terms directly into the GLV equations. In general, we expect the effects of HOIs to alter, but not qualitatively reverse community-level amplification or dampening. This is because HOI's are of order $\mathcal{O}(C^3)$, and thus much weaker than the pairwise interactions (which are $\mathcal{O}(C^2)$) when biomass is relatively low at the early stages of community growth we consider here. Specific structures of HOI's would be needed to override the qualitative (net positive, leading to amplification, or net negative, leading to dampening) effects of pairwise interactions on community-level respiration. For example, if pairwise interactions are on average facilitatory (increasing biomass growth across the community) the sum of all HOIs would need to sum to a negative value to override the effects of the pairwise interactions. Nevertheless, future work focusing on HOI's will likely yield a more nuanced and accurate understanding of the effects of species interactions on microbial community functioning and the dampening or amplification phenomenon.

**Effects of indirect interactions.** The interaction coefficients in the GLV model (Eq 6) represent direct pairwise effects. Indirect effects (competition or facilitatory) between a pair of species can emerge in such communities, through interaction chains. The simplest example involves three species (say, 1, 2, and 3), where a species 1 affects 3 through 2 while the direct interaction coefficient between 1 and 3 is zero. In the case of facilitatory interactions, this could arise, for example, from species 1 producing a metabolic by-product that 2 utilises, and 2 in turn produces a by-product that 3 utilises. In the case of competitive, interactions, a particularly important type of indirect interaction structure is intransitive competition, where strictly hierarchical rock-paper-scissors type orderings of pairwise competition coefficients can stabilise three or more species' biomass dynamics [30,32]. In cases where communities are dominated by indirect interactions, we expect our results about dampening (if $\bar{a} < 0$, when competitive interactions dominate) or amplification (if $\bar{a} > 0$, when facilitatory interactions dominate) of community-level respiration to remain qualitatively unchanged, because the net feedback between biomass and growth rates will still be negative or positive, respectively. For example, intransitive competition enhances coexistence by reducing the abundance of dominant competitors, which is still overall a negative feedback (which would cause dampening of biomass growth). To demonstrate this explicitly, we numerically simulated communities with varying levels of intransitive positive as well as negative interactions. To this end, we randomly generated communities (setting $N = 5$) with different strengths of intransitive interactions and average interaction strength across the community. We then both directly simulated these communities over time using the GLV model (Eq 6) and generated predictions of thermal sensitivity from our theory (Main text Eq 3). Specifically, for each such community, we first generated a $5 \times 5$ random interaction matrix by drawing from a uniform random distribution with mean $\bar{a}$ and range 1. We then set the strength of the intransitive interaction loop (i.e. $a_{12}, a_{23}, \ldots, a_{51}$). We then add a correction factor to all non-intransitive interactions to maintain the desired average interaction strength. This factor was determined by the deviation of total interaction strength from the desired value divided by the number of non-intransitive interactions in the community. The results (Supplementary Figure 1) show that varying the level of intransitive interactions does not affect the qualitative effect of altering the average interaction strength across the community or the predicted thermal or simulated thermal sensitivity values. To illustrate the qualitative robustness of our results, here we modeled communities that were either strongly competitive or facilitatory (bottom black vs top red lines in Supplementary Figure 1), spanning a range of intransitivity (the x-axis of the figure). In regimes where highly intransitive interaction structures are combined with weak overall (average) competition, which have been shown to yield a positive diversity-function relationship [32], would be expected to yield weak amplification of respiration at the community level, or switching between amplification and dampening if the weakly-competitive community structure changes during assembly and subsequent turnover.

**Relation to other methods** Though the derivation of the method we use to infer interactions here is new it has similarities with previously used methods in terms of the overall approach and types and amounts of data used. In this section we highlight similar methods of interaction inference use previously.

The most similar approach is the relative productivity method [33, 43] which is based on the productivity (measured as either biomass accumulation or respiratory rate) of whole communities when compared to the sum of their productivity in monoculture. This is based on the idea that if interactions are neutral then the total community productivity should simply be the sum of the individual strain productivity in monoculture. If they are competitive then productivity will be reduced relative to the neutral case and vice versa. The difference between this metric and ours is that we consider the change in growth at the pairwise level (allowing us to get individual estimates of each pairwise interaction) whereas the relative productivity method measures interactions across the whole community. In this way our metric can be thought of the pairwise version of this measure.

Our approach based on the change in growth rates in monoculture and in pairs is also similar to the relative-yield method[44]. This method compares the total biomass (i.e. carrying capacity) reached by species pairs when grown together verses in isolation. As with the relative productivity method the nature of the interaction is inferred by comparing the yield to the non-interacting case where final paired biomass is simply the sum of the individual carrying capacities.

### Supplementary Figures and Tables

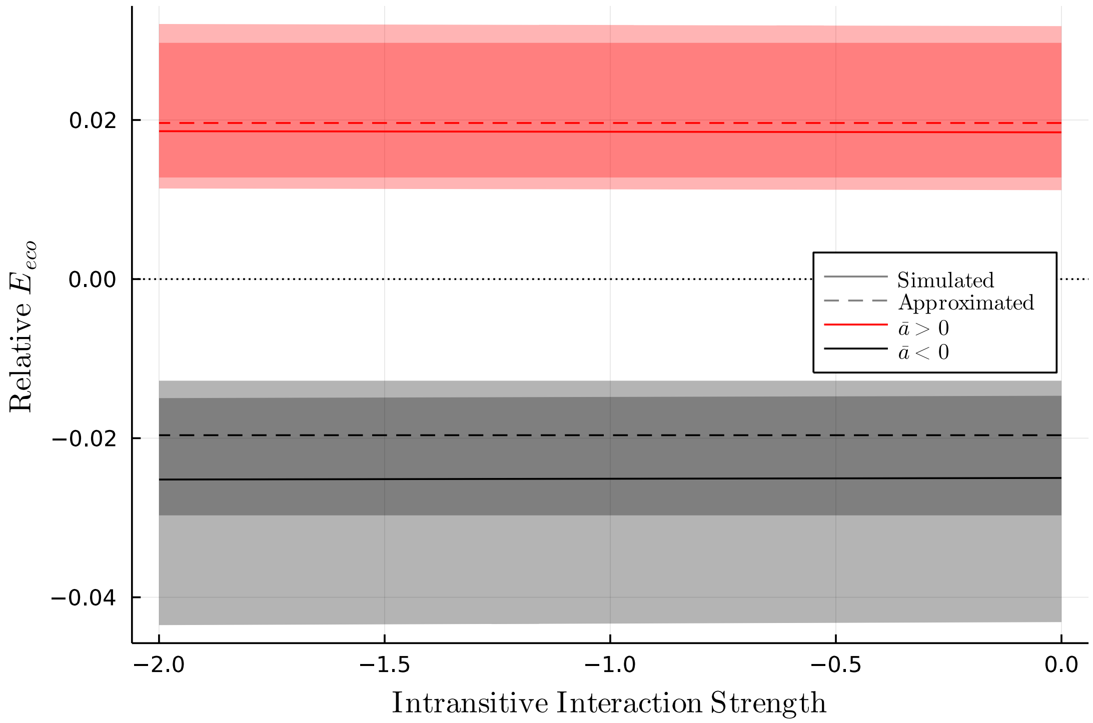


Figure 1: The inclusion of intransitive interaction loops does not affect the predicted thermal sensitivity or the qualitative effect of altering average interaction strength. Lines show the simulated (solid) and approximated (dashed) thermal sensitivity values over differing levels of intransitive interaction strength and average interaction strength (red and black). These simulation results were obtained by numerically integrating the Lotka-Volterra dynamics generated with the same parameters as Fig 1b (see Methods) but with interspecific interactions drawn from a normal distribution such that $a_{ij} \sim \mathcal{N}(\mu_a, 1/N)$ where $\mu_a$ is $0.02$, $0.0$ or $-0.02$ for the facilitatory, neutral and competitive cases respectively (same as the values used to calculate the predicted $E$ values based on the analytical approximation presented in the main text). $E$ values were then obtained by taking the slope of a linear model of $\log(R_{eco})$ vs $\left(\frac{1}{kT} - \frac{1}{KT_{ref}}\right)$ fitted to the simulation data. Shaded areas represent 5-95 percentile bounds calculated across replicates.

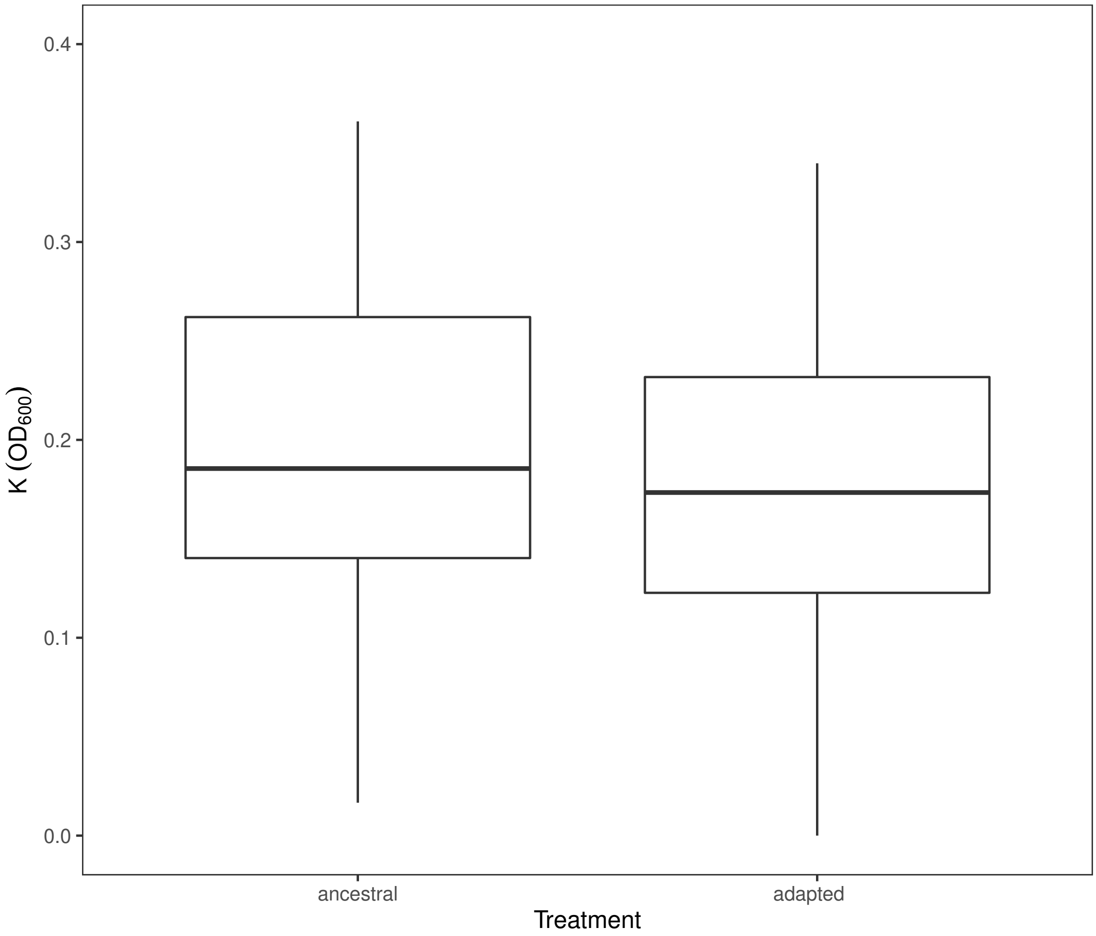


Figure 2: Carrying capacities for single bacterial strains (5 strains, n = 6 technical replicates or each strain) before and after co-adaptation (treatment). Box plots depict the median (centre line) and the first and third quartiles (lower and upper bounds). Whiskers extend to 1.5 times the inter-quartile range (the distance between the first and third quartiles).No significant difference was found between carrying capacities of ancestral and adapted strains (see Table 5 for details of one-sided ANOVA test).

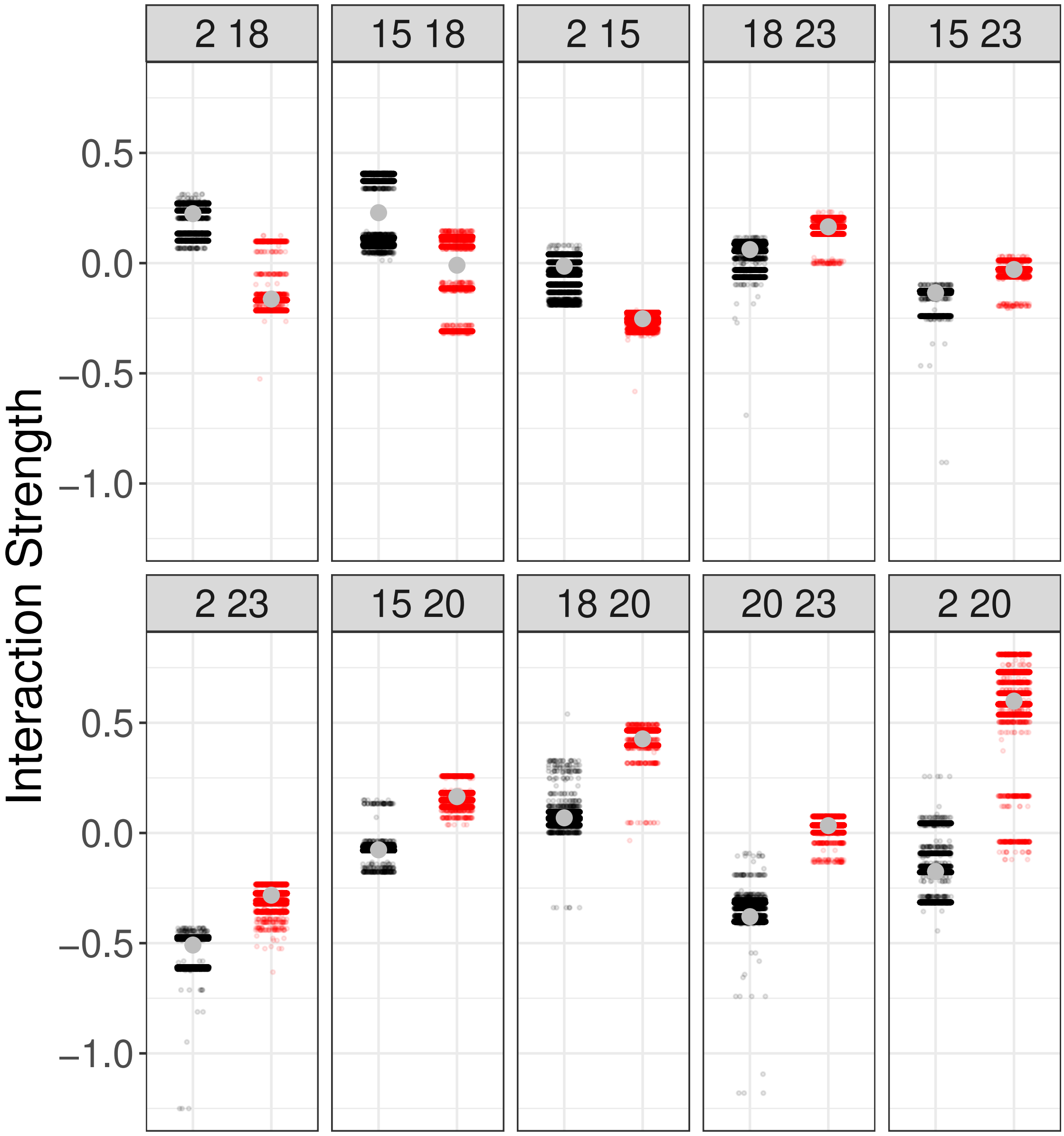


Figure 3: **Pairwise Interaction estimates in the microbial communities** Each panel shows the estimates of interaction strengths between pairs of strains from de novo (black) versus adapted (red) communities. Each point is a single bootstrapped estimate with the mean across all estimates shown by a grey dot. Most (7/10) of the pairs show a shift towards significantly more positive interaction coefficients following longer-term assembly (see Fig 3b). See methods for details on interaction estimation.

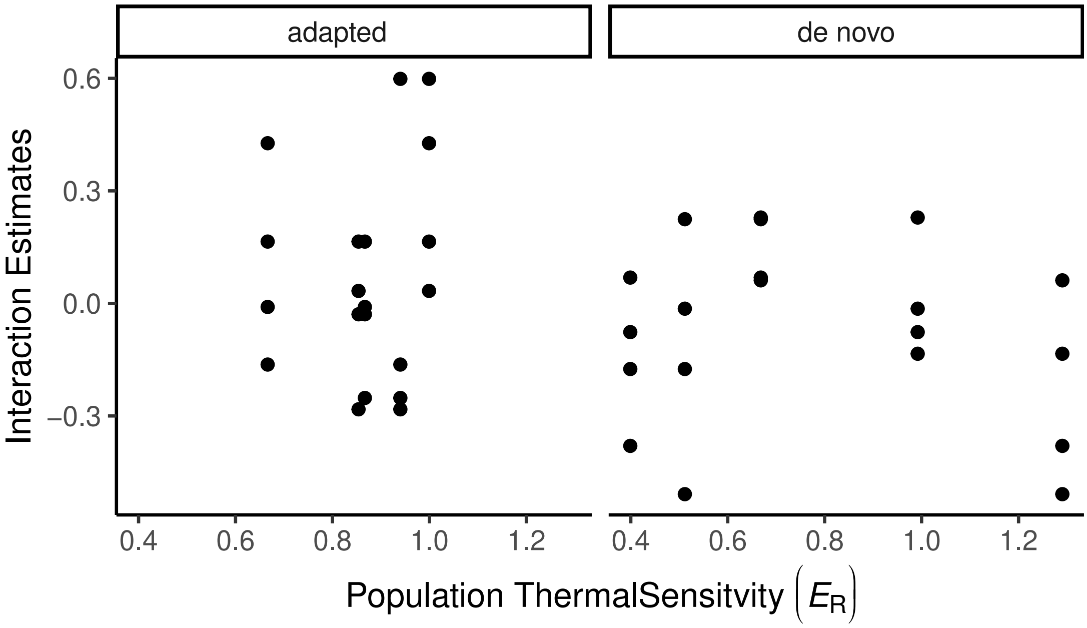


Figure 4: There is no clear relationship between the interaction estimates and population level thermal sensitivity of respiration either before or after assembly. This indicates that the increase in the thermal sensitivity of community respiration cannot be explained by an increase in the tendency for interactions to positively affect populations with high thermal sensitivity.

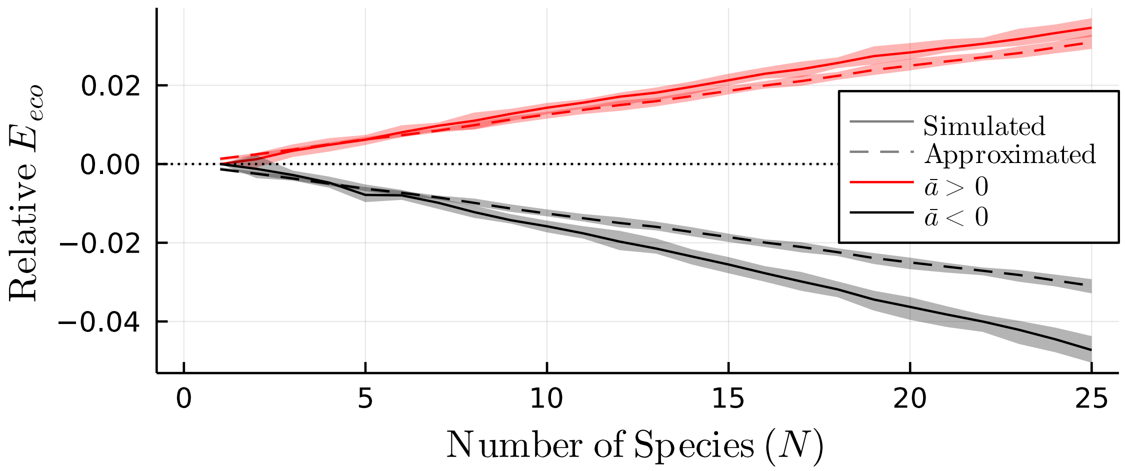


Figure 5: The analytical approximation of thermal sensitivity $E_{eco}$ gives the same qualitative result as numerical simulations using the generalised Lotka-Volterra over varying system sizes. Lines show the simulated (solid) and approximated (dashed) $E_{eco}$ values relative to the neutral interaction case ($\bar{a} = 0$) for facilitatory (red) and competitive (black) interactions. a) The approximate and simulated $E_{eco}$ values show the same sign of change, with both lines remaining on the same side of the horizontal line at $y = 0$ across differing system sizes. The Increase in amplification with increasing system size arises due to increase in total interaction strength experienced by each population as system size increases. Simulations were carried out with the same methods as in Supplementary Figure 1

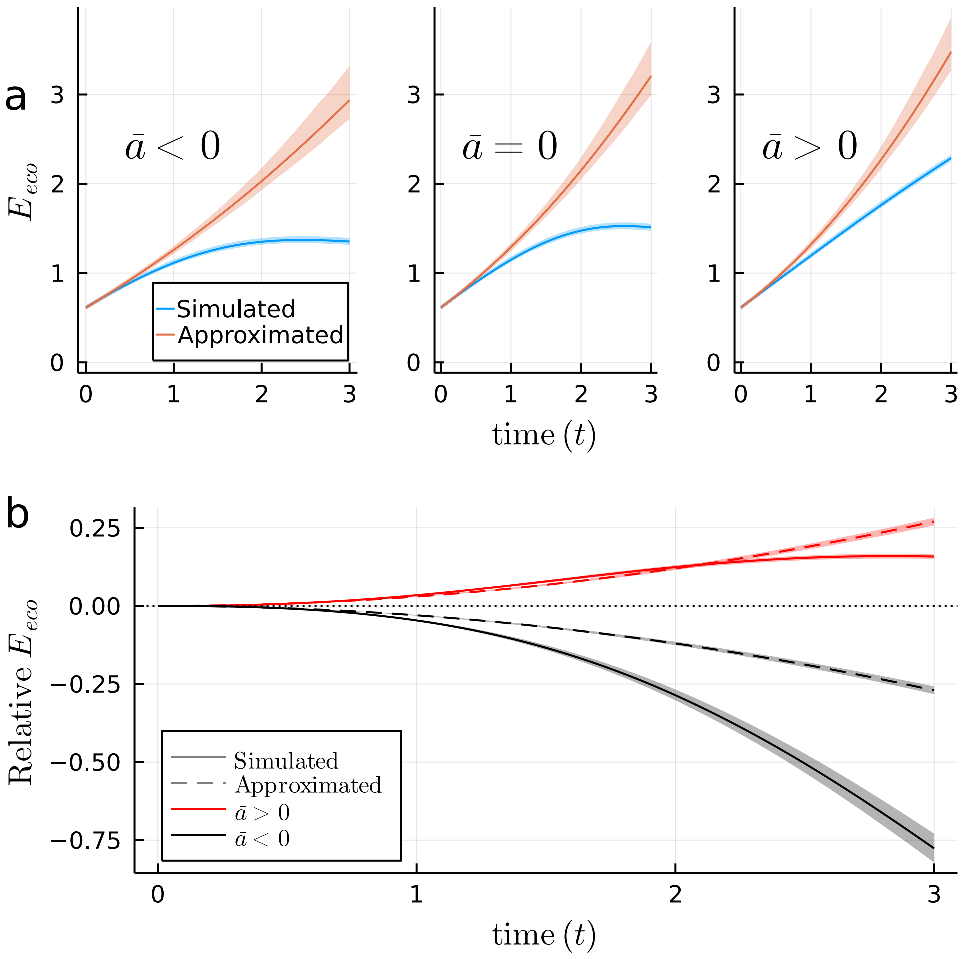


Figure 6: Comparison of the ecosystem thermal sensitivity $E_{eco}$ obtained from mean-field approximation of the the generalised Lotka-Volterra model to that obtained from numerical simulation of the model. a) The thermal sensitivity obtained from numerical simulations (blue) follow the same qualitative pattern as those from the analytical approximation (red), increasing in magnitude as interactions move from being competitive to neutral to facilitatory (moving across panels, from left to right. b) The pattern is made clearer when thermal sensitivity relative to neutral interactions for both the facilitatory ($\bar{a} > 0$, red) and competitive ($\bar{a} < 0$, black) cases are plotted over time. In both interaction scenarios, the simulated (solid) and approximated (dashed lines) exhibit the same sign of change in $E_{eco}$, with both lines remaining on the same side of the horizontal line at $y = 0$. The simulation results were obtained by numerically integrating 100 replicate communities with Lotka-Volterra dynamics (Eq 6) generated with the same parameters as Fig 1b (see methods) but with interspecific interactions drawn from a normal distribution such that $a_{ij} \sim \mathcal{N}(\mu_a, 1/N)$ where $\mu_a$ is 0.02, 0.0 or $-0.02$ for the facilitatory, neutral and competitive cases respectively (same as the values used to calculate the $E$ values based on the analytical approximation presented in the main text). $E$ values were then obtained by taking the slope of a a linear model of $\log(R_{eco})$ vs $\left(\frac{1}{kT} - \frac{1}{KT_{ref}}\right)$ fit to simulation data. Shaded areas in all panels represent 5-95 percentile bounds calculated from the 100 replicates.

| *Model name* | Remove treatment effect on | d.f. | AIC | BIC | Loglik | Test | L.ratio | p-value |
|---|---|---|---|---|---|---|---|---|
| *Nlme.full* | Full model | 13 | 538.83 | 589.95 | -256.41 | | | |
| *Resl.mix1* | ln(r(Tc)) | 12 | 537.37 | 584.55 | -256.68 | 1 vs 2 | 0.54 | 0.46 |
| *Resl.mix2* | ln(r(Tc)+Ea | 11 | 535.66 | 578.91 | -256.83 | 2 vs 3 | 0.29 | 0.59 |
| *Resl.mix3* | ln(r(Tc)+Ea+Eh | 10 | 533.66 | 572.99 | -256.83 | 3 vs 4 | 0.01 | 0.94 |
| ***Resl.mix4*** | All | 9 | 533.06 | 568.45 | -257.53 | 4 vs 5 | 1.39 | 0.24 |
| *Parameters* | ln$(r(T_c))$ | $E_{\mathrm{a}}$ | $E_{\mathrm{h}}$ | $T_{\mathrm{h}}$ | $T_{\mathrm{opt}}$ | | | |
| *Species* | -0.35 | 0.82 | 10.10 | 305.64 | 30.56 | | | |

Table 1: Model selection and parameters of thermal response curves for respiration at the strain level using non-linear mixed effect models. The Sharpe-Schoolfield model was fitted to the respiration rate data quantified over a temperature gradient from 15 °C to 35 °C for all the ancestral and adapted community strains. Models included random effects on each of the parameters by replicate and treatment (ancestral or adapted) as a fixed factor on each parameter. Models were compared via likelihood ratio test using an analysis of variance (ANOVA) where only parameters deemed significant at $p < 0.05$ were retained in the best fitting model (highlighted in bold). The minimum adequate model concludes that there was no difference in the thermal response curves for respiration between the ancestral and adapted strains. The parameter, ln$(r(T_c))$ is the rate of respiration normalized to an arbitrary reference temperature, $T_{\mathrm{c}} = 18°C$, where no low or high temperature inactivation is experienced. $E_{\mathrm{a}}$ is the activation energy (in eV) that characterises the steepness of the slope leading to a thermal optimum. $E_{\mathrm{h}}$ characterizes temperature-induced inactivation of growth above $T_{\mathrm{h}}$, the temperature where half the enzymes are rendered non functional and $T_{\mathrm{opt}}$ is the optimal temperature at which the maximum respiration rate is reached.

| Community in M9 media | | | | | | | | |
|---|---|---|---|---|---|---|---|---|
| *Model name* | Remove treatment effect on | d.f. | AIC | BIC | Loglik | Test | L.ratio | p-value |
| *Nlme.full* | Full model | 10 | 14.71 | 38.78 | 2.64 | | | |
| ***Resl.mix1*** | Ea | 9 | 12.76 | 34.42 | 2.62 | 1 vs 2 | 0.04 | 0.84 |
| *Resl.mix2* | Ea+Th | 8 | 25.23 | 44.48 | -4.61 | 2 vs 3 | 14.47 | <0.001 |
| *Resl.mix3* | ln(r(Tc)+Ea | 8 | 31.58 | 50.83 | -7.79 | 2 vs 4 | 20.82 | <0.001 |
| *Parameters* | $\ln(r(T_c))$ | $E_a$ | $E_h$ | $T_h$ | | | | |
| *De-novo* | -0.34 | 1.04 | 3.71 | 306.8 | | | | |
| *Adapted* | 0.15 | 1.04 | 2.20 | 303.41 | | | | |
| *Community in spent media* | | | | | | | | |
| *Model name* | Remove treatment effect on | d.f. | AIC | BIC | Loglik | Test | L.ratio | p-value |
| *Nlme.full* | Full model | 10 | 20.46 | 44.53 | -0.23 | | | |
| *Resl.mix1* | Eh | 9 | 18.53 | 40.19 | -0.26 | 1 vs 2 | 0.07 | 0.80 |
| ***Resl.mix2*** | ln(r(Tc)+Eh | 8 | 16.90 | 36.15 | -0.45 | 2 vs 3 | 0.37 | 0.54 |
| *Resl.mix3* | ln(r(Tc)+Ea+Eh | 7 | 28.77 | 45.62 | -7.39 | 3 vs 4 | 13.87 | <0.001 |
| *Resl.mix4* | ln(r(Tc)+Eh+Th | 7 | 37.73 | 54.57 | -11.86 | 3 vs 5 | 22.83 | <0.001 |
| *Parameters* | $\ln(r(T_c))$ | $E_a$ | $E_h$ | $T_h$ | | | | |
| *De-novo* | 0.46 | 0.63 | 2.43 | 308.79 | | | | |
| *Adapted* | 0.46 | 1.40 | 2.43 | 299.31 | | | | |

Table 2: Model selection and parameters of thermal response curves for respiration at community level using non-linear mixed effect models. The Sharpe-Schoolfield model was fitted to the respiration rate data quantified over a temperature gradient from 15 °C to 35 °C for all de novo and adapted communities in two different media, M9+glucose and ‘spent media’. Models included random effects on each of the parameters by replicate and treatment (de novo or adapted) as a fixed factor on each parameter. Models were compared via likelihood ratio test using an analysis of variance (ANOVA) where only parameters deemed significant at $p < 0.05$ were retained in the best fitting model (highlighted in bold). The minimum adequate model concludes that there was no difference in the activation energy (Ea) between the de novo and adapted community in M9+glucose media while in the ‘spent media’ we found differences in the Ea between treatments. The parameter, $\ln(r(T_c))$ is the rate of respiration normalized to an arbitrary reference temperature, Tc=18 °C, where no low or high temperature inactivation is experienced. $E_a$ is the activation energy (in eV) that characterises the steepness of the slope leading to a thermal optimum. $E_h$ characterizes temperature-induced inactivation of growth above $T_h$, the temperature where half the enzymes are rendered non functional.

| *Community in M9 media* | | | | | | | | |
|---|---|---|---|---|---|---|---|---|
| *Model name* | Remove treatment effect on | d.f. | AIC | BIC | Loglik | Test | L.ratio | p-value |
| *Nlme.full* | Full model | 10 | 43.95 | 68.01 | -11.97 | | | |
| *Resl.mix1* | Ea+Th | 9 | 41.50 | 63.16 | -11.75 | 1 vs 2 | 0.45 | 0.50 |
| ***Resl.mix2*** | Ea | 8 | 40.07 | 59.32 | -12.04 | 2 vs 3 | 0.57 | 0.45 |
| *Resl.mix3* | ln(r(Tc)+Ea+Th | 7 | 42.68 | 59.53 | -14.34 | 3 vs 4 | 4.61 | 0.03 |
| *Parameters* | $\ln(r(T_c))$ | $E_a$ | $E_h$ | $T_h$ | | | | |
| *De-novo* | -23.55 | 0.60 | 4.17 | 307.40 | | | | |
| *Adapted* | -22.96 | 0.60 | 1.40 | 307.40 | | | | |
| *Community in spent media* | | | | | | | | |
| *Model name* | Remove treatment effect on | d.f. | AIC | BIC | Loglik | Test | L.ratio | p-value |
| *Nlme.full* | Full model | 10 | 157.62 | 181.69 | -68.81 | | | |
| *Resl.mix1* | Th | 9 | 155.62 | 177.28 | -68.81 | 1 vs 2 | 0.00 | 0.98 |
| *Resl.mix2* | Eh+Th | 8 | 154.09 | 173.32 | -69.04 | 2 vs 3 | 0.45 | 0.50 |
| *Resl.mix3* | ln(r(Tc)+Eh+Th | 7 | 152.17 | 169.02 | -69.09 | 3 vs 4 | 0.10 | 0.75 |
| ***Resl.mix4*** | ln(r(Tc)+Ea+Eh+Th | 6 | 150.97 | 165.41 | -69.49 | 4 vs 5 | 0.80 | 0.37 |
| *Parameters* | $\ln(r(T_c))$ | $E_a$ | $E_h$ | $T_h$ | | | | |
| *De-novo* | -22.71 | 0.47 | 2.30 | 306.53 | | | | |
| *Adapted* | -22.71 | 0.47 | 2.30 | 306.53 | | | | |

Table 3: Model selection and parameters of thermal response curves for respiration per cell at community level using non-linear mixed effect models. The Sharpe-Schoofield model was fitted to the respiration rate data quantified over a temperature gradient from 15ºC to 35ºC for all de novo assembled and adapted communities in two different media, M9+glucose and 'spent media'. Models included random effects on each of the parameters by replicate and treatment (de novo or adapted) as a fixed factor on each parameter. Models were compared via likelihood ratio test using an analysis of variance (ANOVA) where only parameters deemed significant at $p < 0.05$ were retained in the best fitting model (highlighted in bold). The minimum adequate model concludes that there was no difference in the activation energy (Ea) between the de novo and adapted community in both M9+glucose media and the 'spent media'. The parameter, $\ln(r(T_c))$ is the rate of respiration normalized to an arbitrary reference temperature, Tc=18 °C, where no low or high temperature inactivation is experienced. $E_a$ is the activation energy (in eV) that characterises the steepness of the slope leading to a thermal optimum. $E_h$ characterizes temperature-induced inactivation of growth above $T_h$, the temperature where half the enzymes are rendered non functional.

| *Community in M9 media* | | | | | |
|---|---|---|---|---|---|
| *Model* | d.f. | AIC | Loglik | Chisq | p-value |
| *Random effects structure* | | | | | |
| *Random = ∼1/Id* | | | | | |
| *Fixed effects structure* | | | | | |
| *1.ln OD ∼ InvT +1* | 4 | 31.88 | -11.94 | - | - |
| *2.ln OD ∼ InvT +Treatment* | 5 | 25.12 | -7.56 | 8.76 | <0.01 |
| *3.ln OD ∼ InvT *Treatment* | 6 | 26.81 | -7.40 | 0.31 | 0.58 |

| *Community in spent media* | | | | | |
|---|---|---|---|---|---|
| *Model* | d.f. | AIC | Loglik | Chisq | p-value |
| *Random effects structure* | | | | | |
| *Random = ∼1/Id* | | | | | |
| *Fixed effects structure* | | | | | |
| *1.ln OD ∼ InvT +1* | 44 | 38.86 | -15.43 | - | - |
| *2.ln OD ∼ InvT +Treatment* | 5 | 40.85 | -15.43 | 0.005 | 0.94 |
| *3.ln OD ∼ InvT *Treatment* | 6 | 30.52 | -9.26 | 12.33 | <0.001 |

Table 4: Model selection and parameters of thermal response curves for total biomass using linear mixed effect models. The Arrhenius equation (Eq 11) was fitted to the biomass data quantified over a temperature gradient from 15 °C to 25 °C for all de novo assembled and adapted communities in two different media, M9+glucose and 'spent media'. Models included a random effect on the intercept by replicate and treatment (de novo or adapted as a fixed factor on both the slope and intercept. Models were compared via likelihood ratio test using an analysis of variance (ANOVA) where only parameters deemed significant at $p < 0.05$ were retained in the best fitting model (highlighted in bold). The minimum adequate model concludes that there was no difference in the activation energy (Ea) between the de novo and stabilised community in the M9+glucose media. However in the 'spent media' the activation energy for total biomass was significantly higher for the stabilised communities compared to the de novo assembled communities. $E_a$ is the activation energy (in eV) that characterises the steepness of the slope leading to a thermal optimum.

| *Strain level* | d.f. | SumSqs | MeanSqs | F | p |
|---|---|---|---|---|---|
| *Treatment* | 1 | 0.004 | 0.004 | 0.33 | 0.57 |
| *Residuals* | 58 | 0.66 | 0.01 | - | - |

Table 5: One-sided analysis of variance (ANOVA) comparing capacities before and after adaptation (treatment) at the strain level (n = 6 technical replicates for each strain and treatment). Analyses reveals no significant effect of adaptation on the carrying capacity at the strain level.

| *Model* | d.f. | SumSqs | MeanSqs | F | p |
|---|---|---|---|---|---|
| *Community Level* | | | | | |
| *Analysis of Variance* | | | | | |
| *K ∼ Treatment * Media* | | | | | |
| *Response: K* | | | | | |
| *Treatment* | 1 | 0.007 | 0.007 | 9.88 | 0.005 |
| *Media* | 1 | 0.004 | 0.004 | 6.83 | 0.02 |
| *Treatment:Media* | 1 | 0.001 | 0.001 | 1.45 | 0.24 |
| *Residuals* | 20 | 0.01 | 0.001 | | |

Table 6: Two-sided analysis of variance (ANOVA) comparing carrying capacities of five taxa before and after adaptation (treatment) at the community level in M9 +glucose and 'spent media' (media) . Analyses reveals a significant effect of adaptation and media on the carrying capacity at the community level and no interaction effect because treatment effect is the same in each media.

| *Isolate ID* | Genus | Family | Phylum | Stream | Tstream | Accession number |
|---|---|---|---|---|---|---|
| *w_Ic161A* | Pseudomonas sp | Pseudomonaceae | Proteobacteria | 1A | 8 | MZ506751 |
| *n_Ic175L* | Chromobacterium sp | Neisseriaceae | Proteobacteria | 5L | 19 | MZ506749 |
| *p_Ic175L* | Iodobacter sp | Neisseriaceae | Proteobacteria | 5L | 19 | MZ506750 |
| *h_Ic174* | Serratia sp | Enterobacteriaceae | Proteobacteria | 4 | 10 | MZ506746 |
| *n_Ic167* | Aeromonas sp | Aeromonadaceae | Proteobacteria | 7 | 11.4 | MZ506748 |
| *d_Ic171* | Mucilaginibacter sp | Sphingobacteriacea | Bacteroidetes | 1 | 17 | MZ506744 |
| *j_Ic165* | Hebaspirillum sp | Oxalobacteraceae | Proteobacteria | 5 | 26.9 | MZ506747 |
| *h_Ic161A* | Janthinobacterium sp | Oxalobacteraceae | Proteobacteria | 1A | 8 | MZ506745 |

Table 7: List of bacterial isolates used in the experiment. Isolate identification codes, taxonomic information, stream ID, average daily water temperature at the time of sampling and GenBank accession numbers are given.
